# CALR frameshift mutations in MPN patient-derived iPS cells accelerate maturation of megakaryocytes

**DOI:** 10.1101/2021.07.05.451130

**Authors:** Kathrin Olschok, Lijuan Han, Marcelo A. S. de Toledo, Janik Böhnke, Martin Graßhoff, Ivan G. Costa, Alexandre Theocharides, Angela Maurer, Herdit M. Schüler, Eva Miriam Buhl, Kristina Pannen, Julian Baumeister, Milena Kalmer, Siddharth Gupta, Peter Boor, Deniz Gezer, Tim H. Brümmendorf, Martin Zenke, Nicolas Chatain, Steffen Koschmieder

## Abstract

Calreticulin (CALR) mutations are driver mutations in myeloproliferative neoplasms (MPNs), leading to activation of the thrombopoietin receptor, and causing abnormal megakaryopoiesis. Here, we generated patient-derived *CALR*ins5- or *CALR*del52-positive induced pluripotent stem (iPS) cells to establish a MPN disease model for molecular and mechanistic studies. We demonstrated myeloperoxidase deficiency in CD15+ granulocytic cells derived from homozygous *CALR*-mutant iPS cells, rescued by repairing the mutation using CRISPR/Cas9. iPS cell-derived megakaryocytes showed characteristics of primary megakaryocytes such as formation of demarcation membrane system and cytoplasmic pro-platelets protrusions. Importantly, *CALR* mutations led to enhanced megakaryopoiesis and accelerated megakaryocytic development in a thrombopoietin-independent manner. Mechanistically, our study identified differentially regulated pathways in mutated vs. unmutated megakaryocytes, such as hypoxia signaling, which represents a potential target for therapeutic intervention. Altogether, we demonstrate key aspects of mutated *CALR-*driven pathogenesis, dependent on its zygosity and found known and novel therapeutic targets, making our model a valuable tool for clinical drug screening in MPNs.

## Introduction

Myeloproliferative neoplasms (MPNs) are a group of clonal hematopoietic disorders including polycythemia vera (PV), essential thrombocythemia (ET), and primary myelofibrosis (PMF), characterized by their excessive increase of granulomonocytic cells, erythroid cells, and/or platelets as well as different degrees of splenomegaly and bone marrow (BM) fibrosis (Campo et al., 2011). Somatic calreticulin (CALR) mutations were discovered in patients with ET and PMF and have been shown to be mutually exclusive with Janus kinase 2 (JAK2) and thrombopoietin (TPO) receptor (*MPL*) mutations (Klampfl et al., 2013; Nangalia et al., 2013). The most common types of *CALR* mutations are 52 bp deletions (del52; type 1) and 5 bp insertions (ins5; type 2), all leading to a +1 frameshift and a common novel C terminus of the CALR mutant protein (Klampfl et al., 2013; Nangalia et al., 2013). Although most *CALR* mutations are heterozygous, homozygous mutations have been observed and shown to be derived from a copy-neutral loss of heterozygosity of chromosome 19p (Klampfl et al., 2013; Stengel et al., 2019; Theocharides et al., 2016).

The oncogenic functions of CALR mutant proteins rely on its binding to the TPO receptor. This leads to the constitutive activation of JAK2 downstream targets such as STAT5, ERK1/2 and AKT, thus causing cellular transformation and abnormal megakaryopoiesis, a hallmark of CALR mutation positive MPNs (Araki et al., 2016; Chachoua et al., 2016; Han et al., 2016; Kollmann et al., 2017; Marty et al., 2016). Megakaryocytes (MKs) are platelet releasing cells, which are known to undergo different stages of maturation (George, 2000; Ru et al., 2015), but the impact of *CALR* mutations on megakaryocytic differentiation and function is still incompletely understood.

It has been reported that patients carrying homozygous *CALR*ins5 exhibit myeloperoxidase (MPO) deficiency as a result of a posttranscriptional mechanism, most likely due to defective CALR chaperone function (Nauseef et al., 1995; Theocharides et al., 2016). Additionally, a *CALR* knock-in mouse model demonstrated that homozygous *CALR*del52 expression resulted in severe thrombocytosis and a myelofibrosis (MF) phenotype (Li et al., 2018). However, the pathogenetic impact of *CALR* mutant zygosity on hematopoietic and megakaryocytic differentiation has not yet been fully addressed.

Several *CALR* mutant disease models have been used in order to uncover its molecular and cellular mechanisms in the pathogenesis of MPN. *CALR* mutant overexpressing cell line models are widely used as the cells can be grown indefinitely in form of clonal populations (Araki et al., 2016; Chachoua et al., 2016; Elf et al., 2016; Han et al., 2016). However, these cell lines are derived from undifferentiated blast populations and, thus, their capacity to differentiate into hematopoietic lineages is limited. MARIMO cells, the only human cell line to date harboring mutated *CALR* (Kollmann et al., 2015), have been reported to carry an additional *NRAS* Q61K mutation which is responsible for transformation, while mutant *CALR* was not essential for these cells (Han et al., 2018). Therefore, the validity of this cell line as a model for *CALR* mutation analysis is questionable. Mouse models are invaluable tools for modelling human disease *in vivo*. Transgenic (Benlabiod et al., 2020; Li et al., 2018; Shide et al., 2019) or retrovirally transduced (Elf et al., 2016; Marty et al., 2016) mutant *CALR* expressing mice have been generated showing MPN-like phenotypes. However, the physiological and genetic differences between species might hamper the translation of human disease phenotypes in the mouse. For these reasons, *CALR* mutant MPN patient samples are desirable for analyzing biological process underlining these diseases. However, the clonal heterogeneity of hematopoietic stem and progenitor cell (HSPC) populations and technical limitations to isolate single clones from patients present major challenges to determine the impact of *CALR* mutant zygosity on clonal composition and diversity in MPN.

Human induced pluripotent stem (iPS) cells are primary cell lines with self-renewal capacity and have the potential to differentiate into any cell type of the three germ layers (Takahashi et al., 2007). Of note, iPS cells can be efficiently differentiated into HSCs allowing *in vitro* hematopoiesis modelling (Sugimura et al., 2017). Patient-derived iPS cells can be further genetically engineered with the CRISPR/Cas9 technology to repair or introduce disease-related mutations in the patient’
ss genetic background. This approach provides a valuable tool to study genotype-phenotype correlations at the clonal level.

Thus, to overcome the aforementioned limitations, we generated MPN patient-specific iPS cell clones carrying homozygous or heterozygous *CALR* mutations or its isogenic unmutated counterpart. *CALR* mutations accelerated megakaryopoiesis independent of TPO, especially in homozygous mutant clones, and repair of the mutation using CRISPR/Cas9 reversed the effect. RNAseq of *CALR*-mutated vs. non-mutated MKs demonstrated increase in hypoxia signaling and leptin expression providing a better understanding of disease biology in ET and PMF and potential novel therapeutic targets.

## Results

### Generation of MPN patient-derived iPS cell clones and repair of homozygous *CALR*ins5 and del52 mutations

iPS cells were established by reprogramming of peripheral blood mononuclear cells (PBMCs) from three MPN patients carrying *CALR*del52, ins5, or del31 mutations (Table S1) and two healthy donors (HD). We obtained 9 homozygous and 55 heterozygous CALRdel52 iPS cell clones derived from a PMF patient, and 74 homozygous and 27 heterozygous *CALR*ins5 clones from a post ET-MF patient (Table S1). Additionally, 64 heterozygous and 5 WT clones were generated from a PMF patient carrying a *CALR* del31 mutation (Table S1). Pluripotency of iPS cells was confirmed by positive staining for OCT3/4 and TRA-1-60 pluripotency markers (Fig. 1 A). Normal karyotype of MPN patient-specific iPS cell clones was shown by GTG-banding (Fig. S1 A). Endogenous expression of mutant *CALR* was confirmed by *CALR* allele-specific RT-qPCR and immunofluorescence (Fig. 1, B and C).

**Figure 1.**
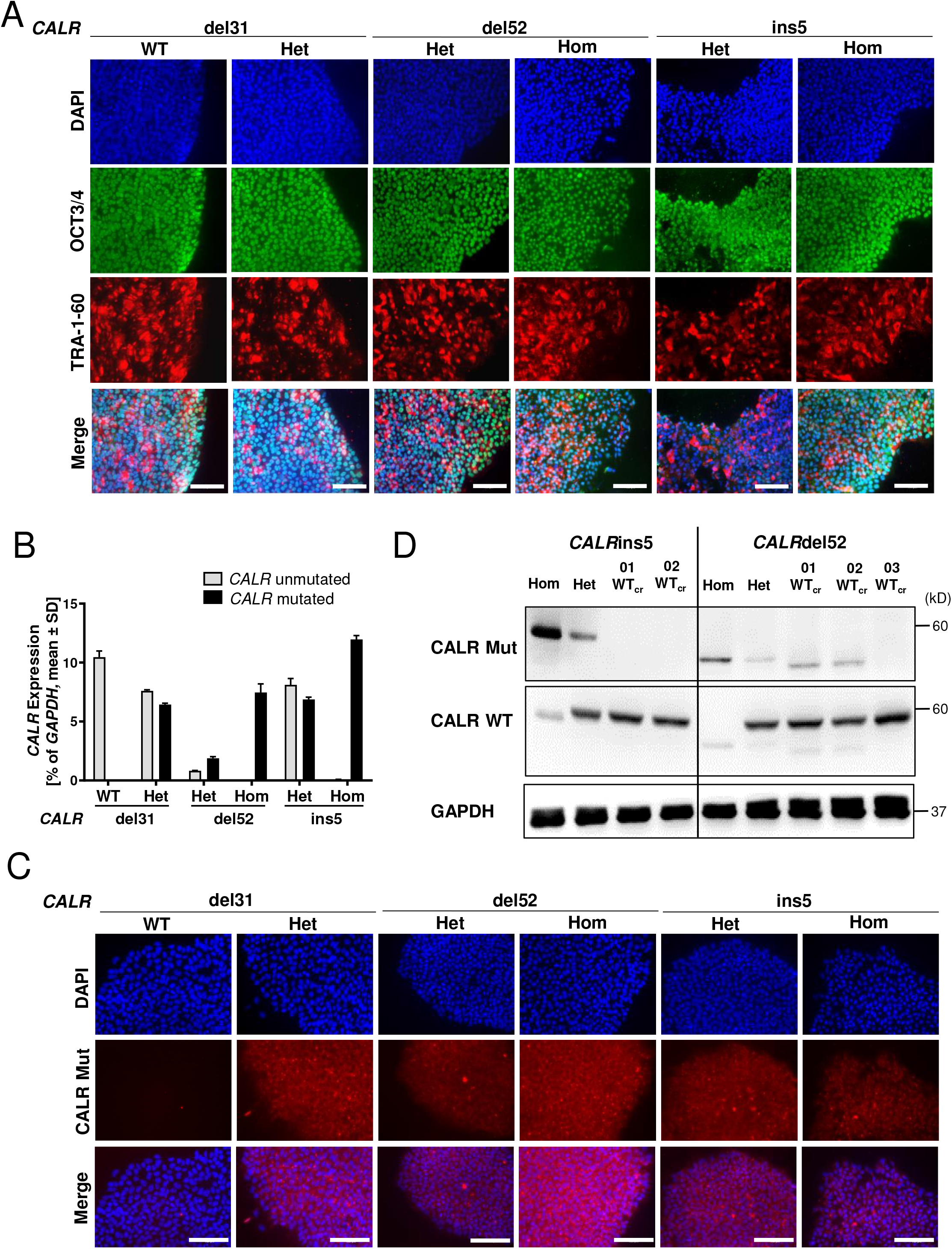
MPN-related iPS cell clones exhibit pluripotency state and harbor *CALR* mutations. **(A)** Pluripotency assessment of indicated iPS cell clones by immunofluorescence. Merge represents overlay of DAPI (blue), OCT3/4 (green) and TRA-1-60 (red). Scale bar, 100 µm. **(B)** *CALR* mRNA expression of indicated iPS cell clones was assessed by *CALR* mutant allele-specific RT-qPCR. Gene expression is depicted as percentage to *GAPDH*. Mean value ± SD of representative clones with indicated *CALR* genotypes. **(C)** Mutant CALR expression (red) was confirmed by immunofluorescence staining with a CALR mutant specific antibody. Unmutated CALR iPS cells served as negative control. Nuclei were stained with DAPI (blue). Scale bars, 100 μm. **(D)** Representative Western blot analysis of CALR WT and CALR mutant (Mut) protein in iPS cell clones after CRISPR repair of homozygous *CALRins5* and del52 mutations. CALR WT protein was assessed on the same membrane as CALR mutant without (w/o) stripping of the CALR Mut antibody explaining residual CALR mutant bands. GAPDH was used as loading control.

Reprogramming of *CALR*ins5 and del52 PBMCs only resulted in homozygous and heterozygous clones but not *CALR* unmutated clones, and thus, *CALR*ins5 and *CALR*del52 mutations were repaired by CRISPR/Cas9 editing to obtain unmutated *CALR* iPS cells. Specific guide RNAs and CALR wildtype donor templates with homology arms were designed (Table S2 and S3). Successful correction of *CALR* mutation in homozygous clones was confirmed by PCR and subsequent Sanger sequencing (Fig. S1, B and C). For *CALR*del52 mutation, three CRISPR clones out of 82 screened clones, and for *CALR*ins5, two clones out of 88 showed a homozygous repair, amounting to a CRISPR efficiency of 3.37 % and 2.27 %, respectively. No off-target effects introduced by CRISPR/Cas9 were found (Fig. S1, D and E). Loss of CALR mutant protein was verified by Western blot analysis in the CRISPR-engineered wild-type (WT_cr_) clones (Fig. 1 D). However, clones 01 WT_cr_ and 02 WT_cr_ of the *CALR*del52 CRISPR approach showed unspecific bands when incubating with CALR mutated antibody.. To preclude that truncated versions of CALR protein resulted after the CRISPR repair, we performed standard amplification of the respective genomic region of the *CALR* gene, Sanger sequencing as well as next generation sequencing (NGS), and absence of mutated genomic *CALR* sequence was confirmed.

Furthermore, the presence of additional MPN-related mutations in all generated iPS cell clones was evaluated by NGS using a defined panel of MPN target genes (Kirschner et al., 2018). Besides the *CALR* mutations, no other clinically relevant MPN-related mutation was found in the generated patient-specific and HD control iPS cell clones (Table S4).

### MPO deficiency is detected uniquely in homozygous *CALR*-mutated iPS cell clones and is restored in repaired *CALR* clones

Patient-derived iPS cells harboring the disease causing mutation(s) allow disease modelling in the culture dish. Homozygous *CALRi*ns5 mutations lead to myeloperoxidase (MPO) deficiency in patients (Theocharides et al., 2016), and we recapitulated this feature *in vitro* by differentiating our iPS cells towards hematopoietic cells in a modified embryoid body (EB)-based protocol (Fig. 2 A) (Kovarova and Koller, 2012). We observed MPO deficiency in iPS cell-derived CD15+ cells harboring homozygous *CALR*ins5 or *CALR*del52 mutations. A significantly smaller CD15+cyMPO+ (cytoplasmic MPO) population was found in cells carrying homozygous *CALR*del52 and *CALR*ins5 mutations (Fig. 2, B and C). Meanwhile, relative *MPO* mRNA expression in homozygous *CALR* cells was upregulated (Fig. 2 D), confirming the post-transcriptional defect (Theocharides et al., 2016). Furthermore, a severe reduction of MPO activity in the homozygous *CALR* mutated cells was confirmed by cytochemical staining (Fig. 2 E). Most importantly, MPO activity was rescued by the correction of homozygous *CALR* mutations as shown for the WT_cr_ clones, proving that homozygous mutant CALR is responsible for this defect.

**Figure 2.**
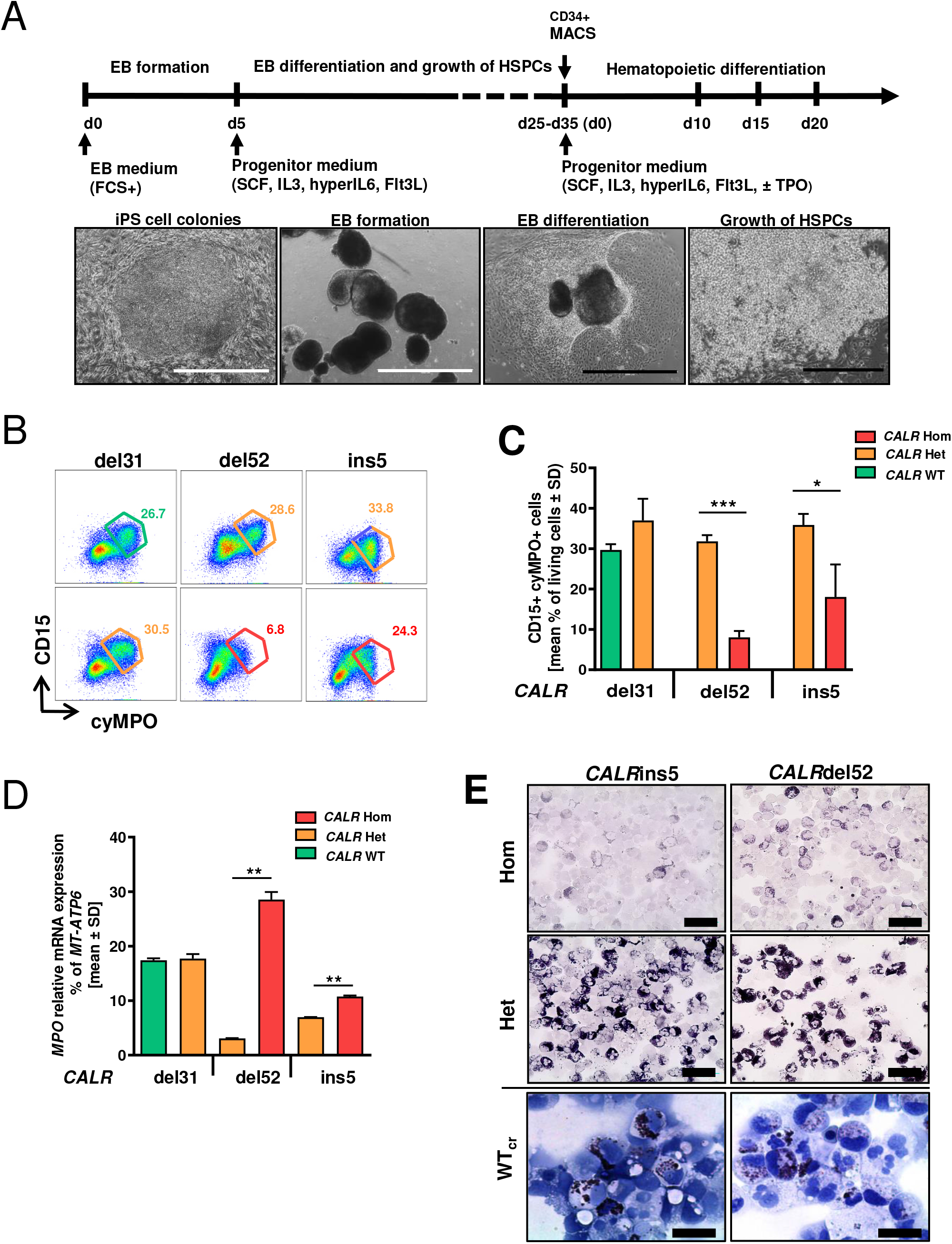
Restoration of MPO activity in iPS cells with repaired *CALRins5* or del52 mutation. **(A)** Schematic representation of a “EB-based” differentiation protocol of iPS cells towards hematopoietic progenitors. Representative images of the differentiation procedure from EB formation to HSPC production of iPS cells are shown. Typical morphology of iPS cell colonies on day 0, embryoid bodies on day 5, mesoderm commitment and differentiated hemogenic endothelial layers on day 8, and HSPC production on day 25. Scale bars white and black, 1000 μm and 400 μm, respectively. **(B)** Flow cytometry gating strategy to identify cyMPO+ neutrophils on day 15 of “EB-based” differentiation. Exemplarily shown for iPS cell-derived hematopoietic cells carrying heterozygous (orange) or homozygous (red) *CALR* mutation or unmutated *CALR* (green). Numbers represent frequencies of CD15+cyMPO+ populations in percent of living single cells. **(C)** Flow cytometry data for cell surface CD15 and intracellular MPO expression of iPS-cell derived HSPCs on day 15 of “EB-based” differentiation. Data of three independent experiments for each *CALR* genotype are shown. \**p*<0.05, \*\*\**p*<0.001. **(D)** Relative *MPO* mRNA expression was confirmed by RT-qPCR. Gene expression is depicted as percentage to *MT-ATP6*. Experiments were performed in triplicates for each *CALR* genotype. Mean value ± SD of representative clones carrying indicated *CALR* genotypes are shown. \*\**p*<0.01. **(E)** MPO functional activity was assessed by cytochemical staining. Representative images of hematopoietic cells harboring indicated *CALR* genotypes are shown. The intensity of the black brown dye indicates the peroxidase activity. Scale bars, 50 μm.

### iPS cells harboring *CALR* mutations give rise to multiple myeloid cell lineages with stronger proliferative capacity

Next, we aimed at getting further mechanistical insights into CALR mutations impacting hematopoietic cells. Therefore, iPS cells that carry the most common type 1 and type 2 *CALR* mutations, *CALR*del52 and *CALR*ins5, respectively, were differentiated towards the myeloid lineage with a modified differentiation protocol designated here as “spin-EB” differentiation (Fig. 3 A) (Liu et al., 2015). From day 8 onwards, hematopoietic cells were released from the EBs into suspension, with an increase of released cells until day 14, as shown in representative images of the differentiation. Homozygous *CALR*ins5- and del52-mutant clones produced a significantly higher number of suspension cells compared to WT_cr_ cells on day 14 (Fig. 3 B). Additionally, cells bearing a heterozygous *CALR*del52 mutation also showed an elevated number of suspension cells than WT_cr_ cells. These data demonstrate that *CALRins5* and *CALRdel52* mutations enhance HSPC proliferation.

**Figure 3:**
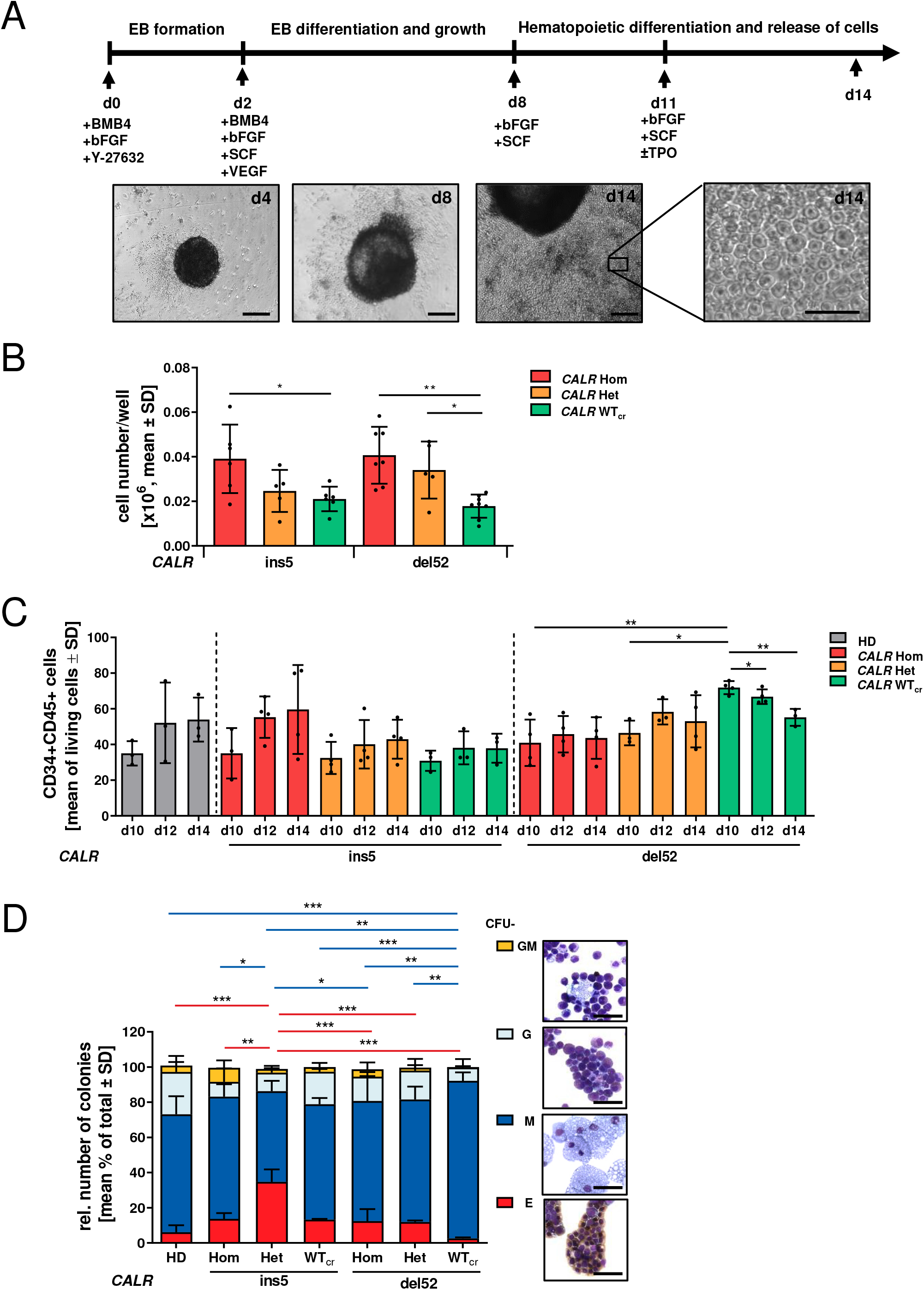
Hematopoietic differentiation potential of iPS cell-derived CD34+ cells. **(A)** Schematic representation of a “spin-EB” protocol to differentiate iPS cells towards HSCs and MKs adapted from Liu et al. (Liu et al., 2015). Cell culture medium was continuously supplemented with cytokines as indicated. On day 14, suspension cells were harvested for further analysis. Representative images of cell culture on indicated days of differentiation are shown. Scale bars, 50 µm. **(B)** Cell number of suspension cells on day 14 of “spin-EB” differentiation calculated for harvested suspension cells/well of 96-well plate. Each data point represents an independent experiment for indicated *CALR* mutation and genotype. \**p*>0.05, ** *p*>0.01. **(C)** Percentage of CD34+CD45+ cells on day 10, day 12 and day 14 of „spin-EB” differentiation analyzed by flow cytometry analysis. Each data point represents an independent experiment for each *CALR* genotype and HD control. \**p*<0.05, \*\**p*<0.01. **(D)** Colony-forming unit (CFU) assay of purified iPS-cell derived CD34+ cells on day 14 of “spin-EB” differentiation. CD34+ cells were isolated by immunogenic bead selection (MACS) and 5,000 cells/ml were seeded in semisolid medium supplemented with cytokines. After 12 days, colonies were characterized and counted. Results of three independent experiments for each *CALR* genotype and HD controls are shown. Red dash and blue dash referred to significant differences in CFU-E and CFU-M, respectively. \**p*>0.05, ** *p*>0.01, \*\*\**p*>0.001. Morphological appearance of different colony types was assessed by cytospin preparation and Diff-Quik staining. Scale bars, 50 µm.

Flow cytometry analysis on day 10, 12 and 14 of suspension cells confirmed generation of CD34+CD45+ HSPCs (Fig. 3 C; S2, A and B). Noteworthy, homozygous and heterozygous *CALR*del52 mutated iPS cells, but not WT_cr_ iPS cells, showed a stable level of HSPCs over the analyzed days. In WT_cr_ cells, a significant reduction of HSPCs over the iPS cell differentiation time was observed accompanied by an overall higher number of HSCs generated on day 10 compared to hetero- and homozygous mutant cells.

To study the myeloid progenitor potential of iPS cell-derived HSCs, we performed a colony-forming-unit assay using CD34+ cells isolated on day 14 of differentiation, previously cultured in the presence of TPO. The heterozygous mutant *CALR*ins5 CD34+ cells produced more colonies compared to WT_cr_ cells (Fig. S2 C). The same trend was observed for the homozygous *CALR*ins5- and *CALR*del52 cells compared to control (*p*=0.0898 and *p*=0.0851, respectively). To examine whether *CALR* mutation impacts on the expansion of CD34+ cells, expression of the proliferative marker *Ki67* was evaluated in RT-qPCR (Figure S2D). We found that HSCs harboring a heterozygous *CALR*ins5 mutation showed higher *Ki67* expression (although not statistically significant, *p*=0.1411) compared to WT_cr_ HSPCs. Together, those data confirmed a stronger amplification of cells in the *CALR*-mutated background, as described above. Identities of CFU-E (erythrocytic cells), CFU-M (macrophages), CFU-G (granulocytes) and CFU-GM (granulocytes/macrophages) were determined by morphological appearance and confirmed by QuikDiff staining (Fig. 3 D). HSPCs from all analyzed clones gave rise to CFU-E, CFU-M, CFU-G and CFU-GM colonies. While *CALR*del52 WT_cr_ HSPCs differentiated into the highest number of CFU-M, they showed less CFU-E. An increased number of erythrocytic colonies were found in *CALR*ins5 heterozygous clones.

In summary, CALR mutants accelerate cell proliferation and *CALR*-mutated iPS cells show constant production of HSPCs.

### *CALR*-mutated iPS cells exhibit enhanced megakaryopoiesis and accelerated maturation in a TPO independent manner

Aberrant megakaryopoiesis is the major hallmark of *CALR*-mutated MPN patients as well as in established mouse models. Therefore, we studied the impact of *CALR*-mutant zygosity on megakaryopoiesis in our patient-specific iPS cell model in our EB differentiation protocol (Fig. 3 A) (Liu et al., 2015). We found that iPS cells carrying homozygous or heterozygous *CALR*ins5 and *CALR*del52 mutation gave rise to significantly more CD41+CD42b+ mature MKs compared to their WT_cr_ counterparts (Fig. 4 A). Moreover, no difference in MK numbers obtained from HD or WT_cr_ iPS cells was observed, demonstrating that the enhanced megakaryopoiesis is due to the *CALR* mutation. Mature megakaryocytes with typical multi-lobular nuclei and slight basophilic cytoplasm were detected in cytospins generated on day 14 of differentiation (Fig. 4 B). Of note, MKs carrying homozygous CALR mutation presented variable sizes, indicative of rapid maturation of aberrant MKs.

**Figure 4.**
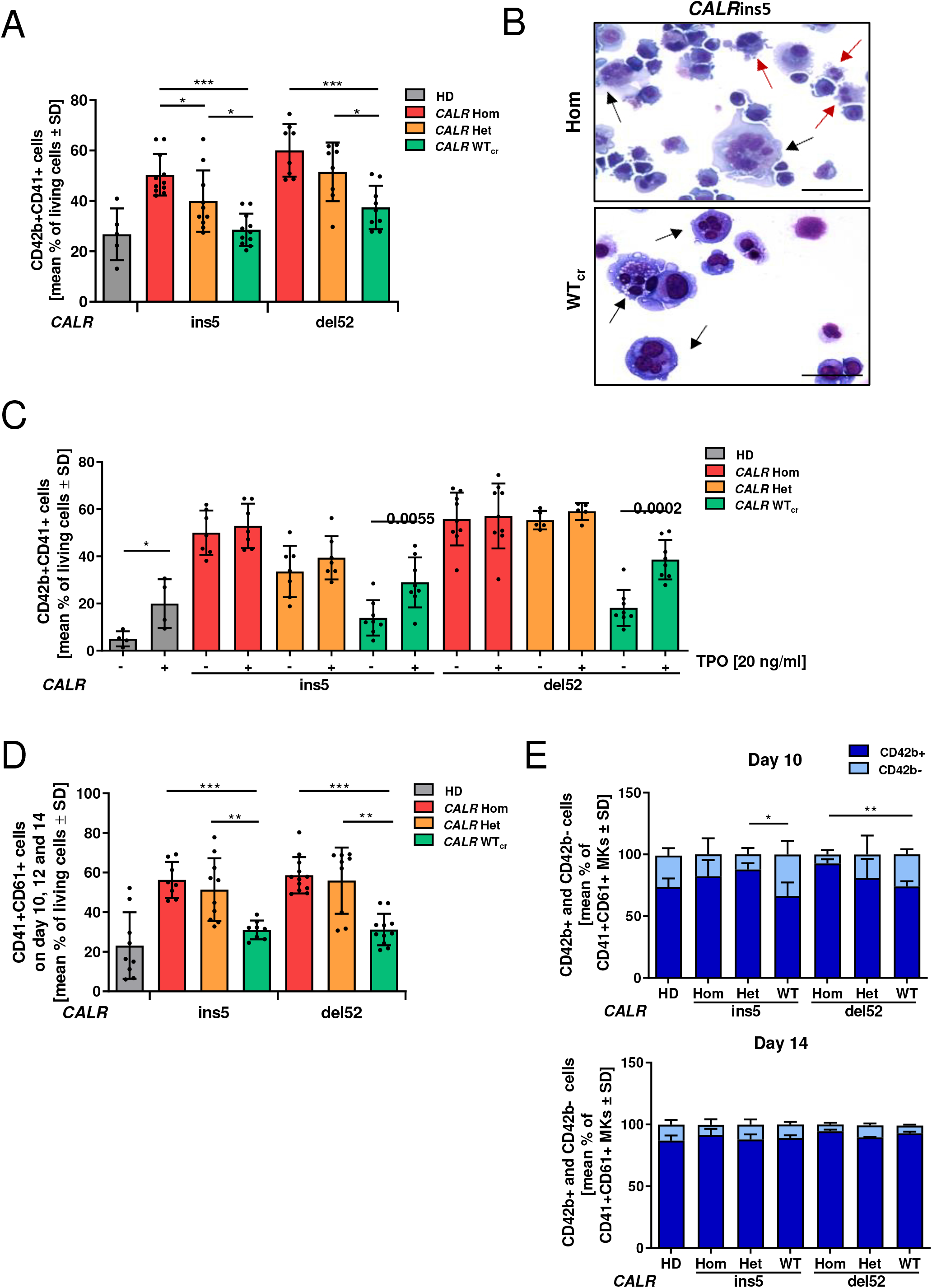
Megakaryocytic differentiation of patient-specific *CALR*ins5 and *CALR*del52 iPS cells. **(A)** Percentage of CD42b+CD41+ matured MKs determined by flow cytometry on day 14 of “spin-EB” differentiation. Numbers of independent experiments performed for each *CALR* genotype and HD control referred to number of data points shown. \**p*<0.05, \*\**p*<0.01, \*\*\**p*<0.001. **(B)** Representative morphology of MKs harvested on day 14 of “spin-EB” differentiation exemplarily shown for *CALRins5* homozygous clone and repaired WT clone (WT_cr_) stained with Diff-Quik solutions after cytospin preparation. Typical MKs and aberrant smaller MKs are indicated with black and red arrows, respectively. Scale bars, 50 µm. **(C)** Impact of TPO on MK development analyzed by flow cytometry analysis on day 14 of “spin-EB” differentiation. Percentage of CD42b+CD41+ MKs is shown for cells treated with (+) or without (-) TPO from day 11 of differentiation onwards. Numbers of independent experiments for each *CALR* genotype and HD control referred to number of data point shown. \**p*<0.05, \*\**p*<0.01, \*\*\**p*<0.001. **(D)** Percentage of CD61+CD41+ immature MKs determined by flow cytometry analysis. Data of day 10, day 12 and day 14 of three independent experiments are combined for each *CALR* genotype and HD controls. \*\**p*<0.01, \*\*\**p*<0.001. **(E)** CD42b+ and CD42b-cells in CD61+CD41+ MKs on day 10 and day 14 of “spin-EB” differentiation analyzed by flow cytometry analysis. Statistical analysis compares number of CD42b+ cells of homozygous clones to corresponding heterozygous and WT_cr_ clones of three independent experiments. \*\**p*<0.01.

Mutant CALR binds to and activates the TPO receptor. To further study the impact of TPO on MK development in our iPS cell model, we differentiated the cells with and without supplementation of TPO (Fig. 4 C). In HD control cells as well as in WT_cr_, we found a significantly lower number of MKs in the absence of TPO. In contrast, the number of iPS cell-derived MKs harboring a *CALR* mutation was equal regardless of *CALR* mutation zygosity and the presence or absence of TPO. Thus, our *CALR*-mutated iPS cell-model recapitulates a key pathological feature of TPO-independent megakaryopoiesis.

By following *CALR*-mutated megakaryopoiesis *in vitro*, we observed a striking increase in immature CD41+CD61+ MKs in comparison to unmutated cells combining flow cytometry data from day 10, 12 and 14 (Fig. 4 D). We further analyzed the portion of mature MKs in the CD41+CD61+ cell population to analyze differences in the MK maturity level along the differentiation. A more in-depth analysis revealed less CD42b+CD41+CD61+ mature MKs from *CALR*del52 WT_cr_ iPS cells when compared to their homozygous counterparts on day 10 of differentiation (Fig. 4 E). The same was observed for the heterozygous *CALR*ins5 clone and with a tendency (*p*=0.1794) also for homozygous *CALR*ins5-mutated MKs. However, on day 14 all genotypes showed the same MK maturity. Altogether, these data demonstrate an accelerated MK maturation process in *CALR*-mutated cells.

MKs arise from progenitors shared with the erythrocytic lineage, the megakaryocyte-erythrocytic progenitors (MEP). Therefore, we analyzed the proportion of erythrocytic cells (CD235a+CD45-) of the day 14 population (Fig. S2 E). HD control iPS cells and iPS cells with *CALR*del52 background originated in an extremely low level of erythrocytic cells. However, *CALR*ins5-mutated cells showed an increase in the erythrocytic population with a remarkable increase in the WT_cr_ cells. These data demonstrate that the repair of the *CALR* mutation in the homozygous *CALR*ins5-mutated cells led to enhanced erythropoiesis, which was not observed in HD control iPS cells or in the WT_cr_ of *CALRdel52* cells. Based on this, we hypothesized that differentiation properties switch upon the repair of the *CALR* mutation from the megakaryocytic to the erythrocytic lineage, at least in the patient-specific background of our *CALR*ins5-repaired clones.

### *CALR* mutation causes upregulation of MK-related genes in iPS cell-derived MKs

MK development is subject to a highly coordinated process of megakaryocyte-specific gene expression and simultaneous prevention of erythrocytic development. To examine whether *CALR*-mutated and -unmutated MKs show differences at the transcriptional level, we analyzed gene expression of typical megakaryocytic genes and related transcription factors. Our clonal approach using iPS cell-derived CALR mutant MKs and their repaired counterparts are especially suitable for the analysis of solely mutated CALR-related changes.

To exclusively analyze the megakaryocytic subpopulation, CD61+ MKs were purified by magnetic activated cell sorting on day 14 of “spin-EB” differentiation. We compared the differential expression signatures of MKs within the group of *CALRins5* MKs and *CALR*del52 MKs in a heat map with unsupervised clustering (Fig. 5A, and B). We found that gene expression profiles of MKs with heterozygous or homozygous *CALR*ins5 mutation built a well-defined cluster compared to unmutated MKs. Of note, heterozygous *CALR*ins5-mutated MKs showed a stronger upregulation of MK-related genes compared to the homozygous *CALR*ins5 mutated MKs. Focusing on genes known to be involved in megakaryocytic development, we found that the expression of MPL was slightly enhanced in *CALR*ins5 heterozygous clones and homozygous clones (*p*=0.0669, Fig. 5 C) compared to WT_cr_ MKs. Moreover, the early transcription factor *FLI1* was highly upregulated in *CALRins5* heterozygous and homozygous MKs compared to WT_cr_ MKs (*p*= 0.0106 and *p*=0.051, respectively). On the other hand, *NFE2*, an essential marker for terminal MK maturation involved in platelet release and expression of *VWF*, was strongly expressed in both homozygous and heterozygous *CALRins5*-mutated MKs (Tijssen and Ghevaert, 2013). Given that homozygous *CALR*ins5-mutated MKs showed an upregulation of mature MK markers (*NFE2* and *VWF*), this suggests enhanced megakaryopoiesis of homozygous mutant cells. In the iPS cell-derived HSPCs, we tested the expression of *Ki67* in MKs, showing equal expression in both mutated *CALR*ins5 and *CALR*del52 and unmutated MKs (Fig. S2 F).

**Figure 5.**
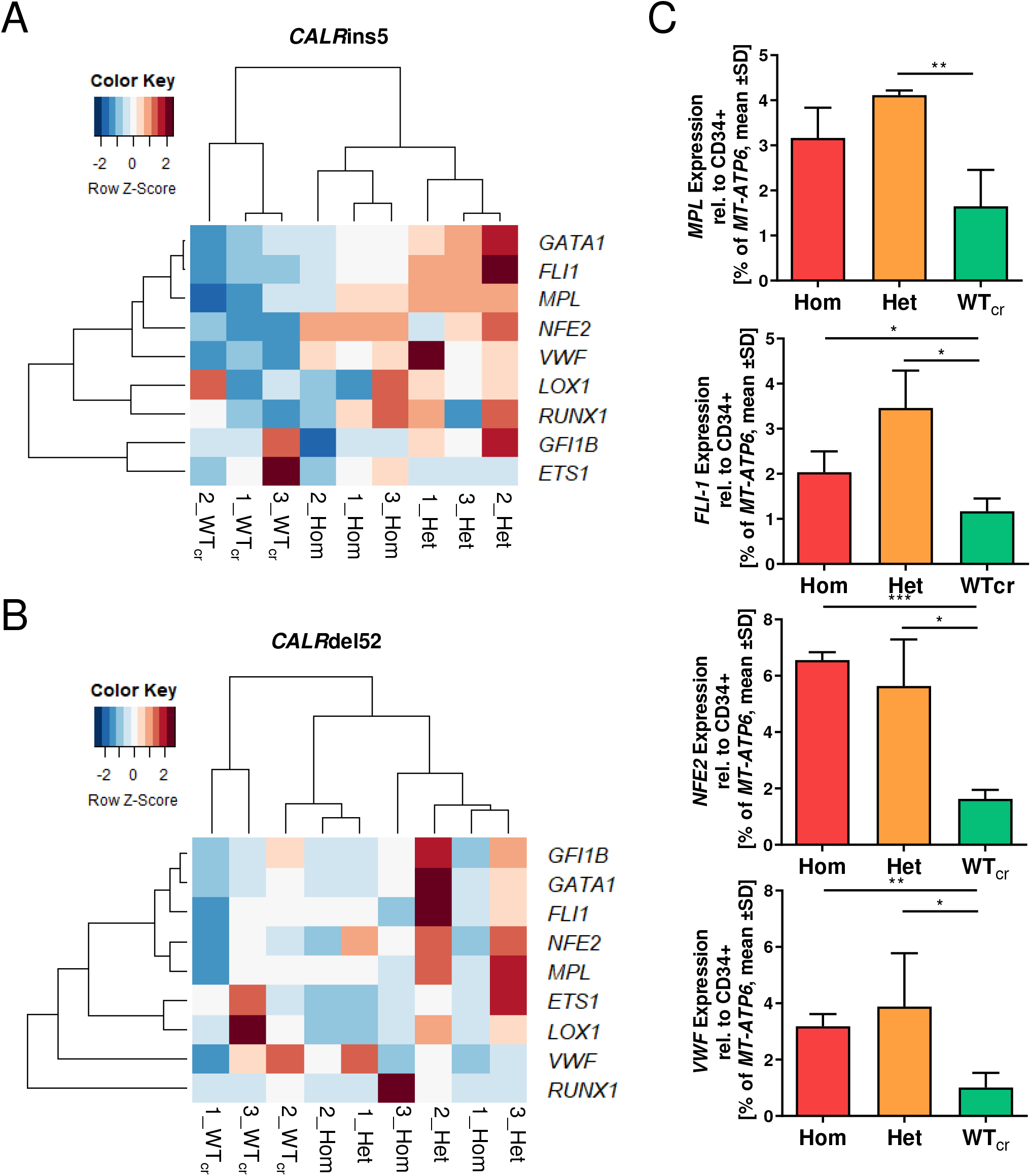
Gene expression profile of iPS cell-derived MKs harboring *CALR* mutations. **(A)** and **(B)** Heat map for unsupervised hierarchical linkage clustering of selected MK-related genes analyzed by RT-qPCR of CD61+ cells. CD61+ cells were isolated by immunogenic bead selection (MACS) on day 14 of “spin-EB” differentiation. Data are normalized to gene expression of purified CD34+ cells of the same experiment. Calculated z-score is shown for individual experiment of indicated *CALR* genotype (ins5 **(A)**, del52 **(B)**, red and blue, high and low expression; respectively). **(C)** Gene expression profile of *MPL, FLI-1, NF-E2*, and *VWF* of CD61+ MKs. Expression was normalized to CD34+ cells from the same experiment. Analysis was performed for three independent experiments performed for each *CALRins5* genotype. Statistical analysis compares the expression of homozygous and heterozygous *CALR*-mutated MKs to WT_cr_ MK. \**p*<0.05, \*\**p*<0.01, \*\*\**p*<0.001.

Unsupervised clustering of MKs with *CALR*del52 mutation did not resultin clear clusters of each zygosity as given for *CALR*ins5-mutated MKs (Fig. 4 B). Nonetheless, gene expression of mutated MKs, both heterozygous and homozygous, clustered closer together than WT_cr_ MKs. In line with the expression data of *CALR*ins5-mutated MKs, the strongest upregulation of MK related genes was found in heterozygous clones, suggesting an underlying mechanism including both mutated and WT protein, which enhances transcription of megakaryocytic genes, likely involving CALR chaperone function.

### *CALR*-mutated MKs exhibit higher granularity

During megakaryopoiesis, MKs increase in size, produce alpha and dense granules, and develop a demarcation membrane system (DMS), the plasma membrane for future platelets (Ru et al., 2015). Terminally differentiated MKs form cytoplasmic protrusions, called pro-platelets, which further mature into platelets with open canalicular systems (OCS) (Escolar and White, 1991).

To study in more detail whether structural features of MK maturation are present in our iPS cell-derived MKs, we captured ultrastructural transmission electron microscopy (TEM) images of CD61+ MKs on day14 of “spin-EB” differentiation for each *CALR*ins5 genotype. TEM images of MKs with different genetic background revealed typical megakaryocytic structures including granules, DMS, pro-platelet protrusions and OCS (Fig. 6 A). Additionally, we determined the area of each MK from the TEM images of each *CALR* genotype (Fig. 6 B). WT_cr_ MKs showed a slight increase (*p*=0.1679) in size compared to homozygous mutant MKs. This supported our previous observation, that some small MKs were found in the population of homozygous mutant MKs in cytospin images (Fig. 4 B). It is reported that *in vitro* generated MKs undergo different stages of maturation, identifiable by the granularity of the cells defined by side scatter (SSC) in flow cytometry (Sim et al., 2017). Therefore, iPS-derived CD42b+CD41+CD61+ MKs were analyzed for their MFI of SSC. We found that the granularity of MKs is significantly enhanced in the homozygous *CALR*ins5 MKs compared to the unmutated counterparts (Fig. 6 C), confirming that homozygous *CALRi*ns5-mutated MKs showed higher level of maturation compared to the unmutated MKs. No differences in the granularity of CALRdel52-mutated MKs was observed, again suggesting that type 1 and type 2 *CALR* mutations may not induce identical functional changes, as is also suggested by differences in their clinical profile.

**Figure 6.**
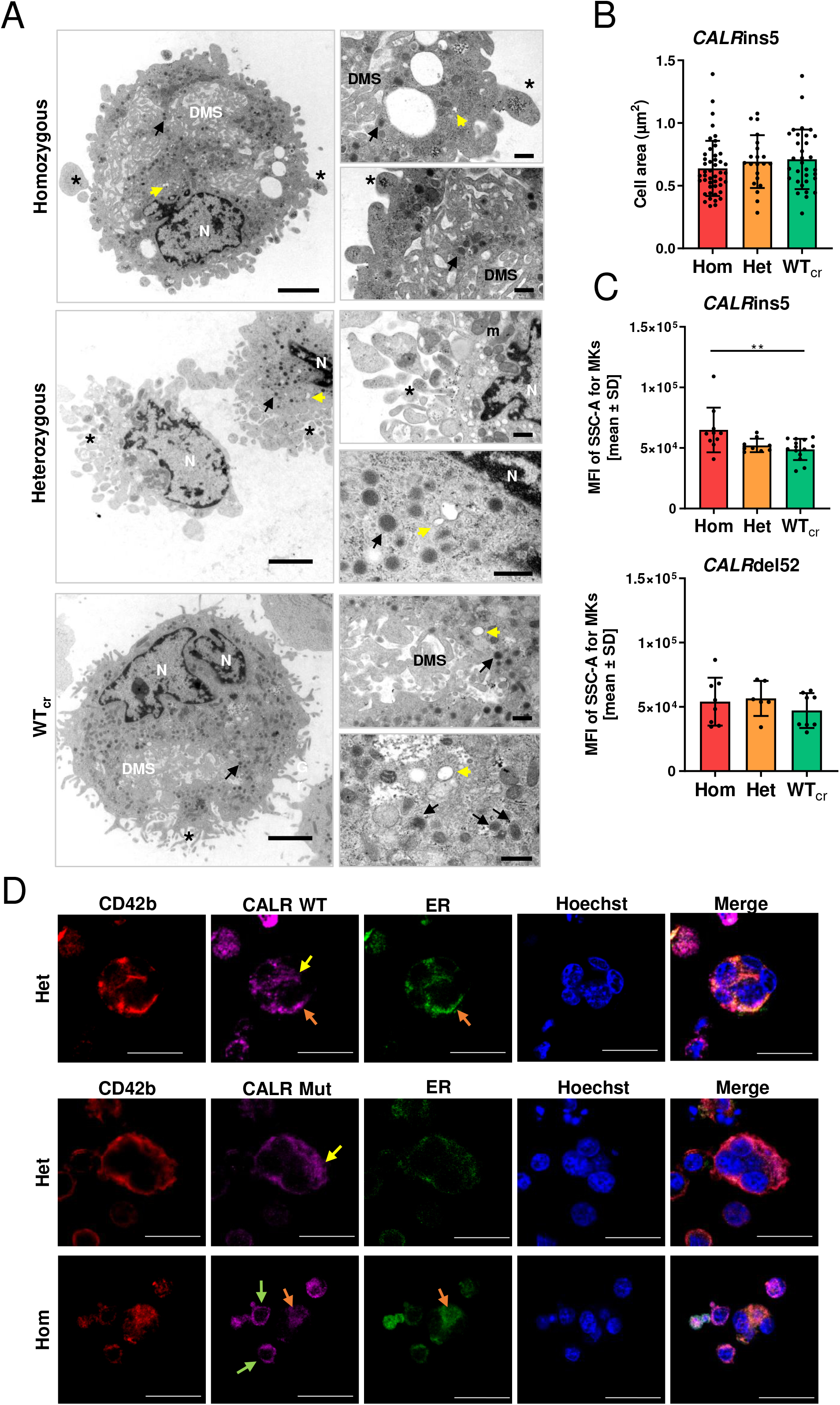
Ultrastructural morphology analysis of iPS cell-derived MKs and CALR distribution in MKs. **(A)** Transmission electron microscopy (TEM) images of iPS cell-derived MKs of indicated *CALR*ins5 genotype to evaluate cell morphology. Representative images are shown. N: nucleus. DMS: Demarcation membrane system. *: pro-platelet protrusions. Black arrow: granules. Yellow arrow: open canalicular system. m: mitochondria. Scale bars, 2.5 µm (large panels) and 500 nm (small panels). **(B)** Calculated cell area of MKs in TEM images for indicated *CALR*ins5 genotypes. Each data point represents a single cell. **(C)** Mean fluorescence intensity (MFI) calculated for the side-scatter (SSC-A) of CD42b+CD41+CD61+ MKs on day 14 of “spin-EB” differentiation. Indicated data points represent independent experiments for each *CALRins5* and del52 genotype. \*\**p*<0.01. **(D)** iPS cell-derived MKs were fixed and stained for the endoplasmic reticulum (ER) and WT CALR or mutated (Mut) CALR after 14 days of differentiation for indicated *CALR*ins5iPS cell clones. To identify MKs, samples were additionally stained for CD42b. Hoechst was added for nuclear staining. Scale bars, 50 µm. Diffuse CALR distribution, clustered localization of CALR at the cell surface, and co-localization of CALR and ER are indicated with yellow, green, and orange arrows, respectively

In the following, we analyzed the distribution of CALR WT and mutant protein in our iPS cell-derived MKs by immunofluorescence and confocal microscopy (Fig. 6 D; S3 A). We were able to identify CD42b positive MKs with pronounced poly-lobulated nuclei in mutated and unmutated MKs, highlighted in z-stacks of homozygous *CALR*ins5 and WT MKs (Fig. S3 A). WT and mutated CALR protein were not differentially distributed in the MKs, and showed a more diffuse distribution in bigger MKs (yellow arrows). However, smaller MKs with less pronounced cytoplasm, showed a clustered localization of mutated CALR at the cell surface (green arrows). In some MKs, we were able to verify a co-localization of CALR protein and the endoplasmic reticulum (ER) (orange arrows).

### *CALR* mutation causes an upregulation of hypoxia related pathways in MKs

Finally, to further unveil differences between RNA expression dependent on the zygosity of the *CALR* mutation, we performed RNA sequencing (RNAseq) analysis of day 14 CD61+ iPS cell-derived MKs. Three independent differentiation experiments were performed using either WT_cr_, heterozygous or homozygous *CALR*ins5-mutated iPS cells. RNAseq data were normalized using TMM normalization (Robinson and Oshlack, 2010) and subsequently voom transformed (Law et al., 2014).

Unsupervised clustering of differentially expressed genes (DEGs) was performed and depicted in a heat map (Fig. 7 A). Samples efficiently clustered according to their *CALR* mutation zygosity, and higher similarity in the gene expression profile of mutated MKs was observed. Unsupervised sub-clustering of the heat map provided cluster 1 with mainly upregulated genes and cluster 2 with downregulated DEGs in homozygous mutant MKs. Gene ontology (GO) analysis of those clusters showed that MKs with homozygous *CALR* mutation upregulated interferon signaling and the response to hypoxia, while ECM organization was downregulated compared to WT_cr_ MKs as exemplarily shown for *FN1* (Fig. 7 B). Of note, GO term including genes related to the ER were found to be affected in heterozygous and homozygous mutant MKs. The observed upregulation of hypoxia signaling was further validated by analysis of Pathway Responsive Genes (PROGENy) (Fig. 7 C) (Schubert et al., 2018). Moreover, the known hypoxia related gene *NDRG1* was significantly upregulated in both heterozygous and homozygous *CALR*ins5-mutated MKs compared to WT. (Fig. 7 D). Interestingly, PROGENy analysis showed that PI3K signaling was significantly downregulated in both heterozygous and homozygous mutant MKs compared to WT, similar to TGFb and EGFR pathways, which were downregulated in the homozygous mutant MKs compared to control MKs. In addition, a high number of DEGs were found by comparing different genotypes (Fig. S5). Genes found to be significantly upregulated in both heterozygous and homozygous mutant MKs compared to WT_cr_ MKs, included *H3C3, CTSF*, and leptin. Partial validation of depicted genes was performed by RT-qPCR (Fig. 7 D; Fig. S5 D).

**Figure 7.**
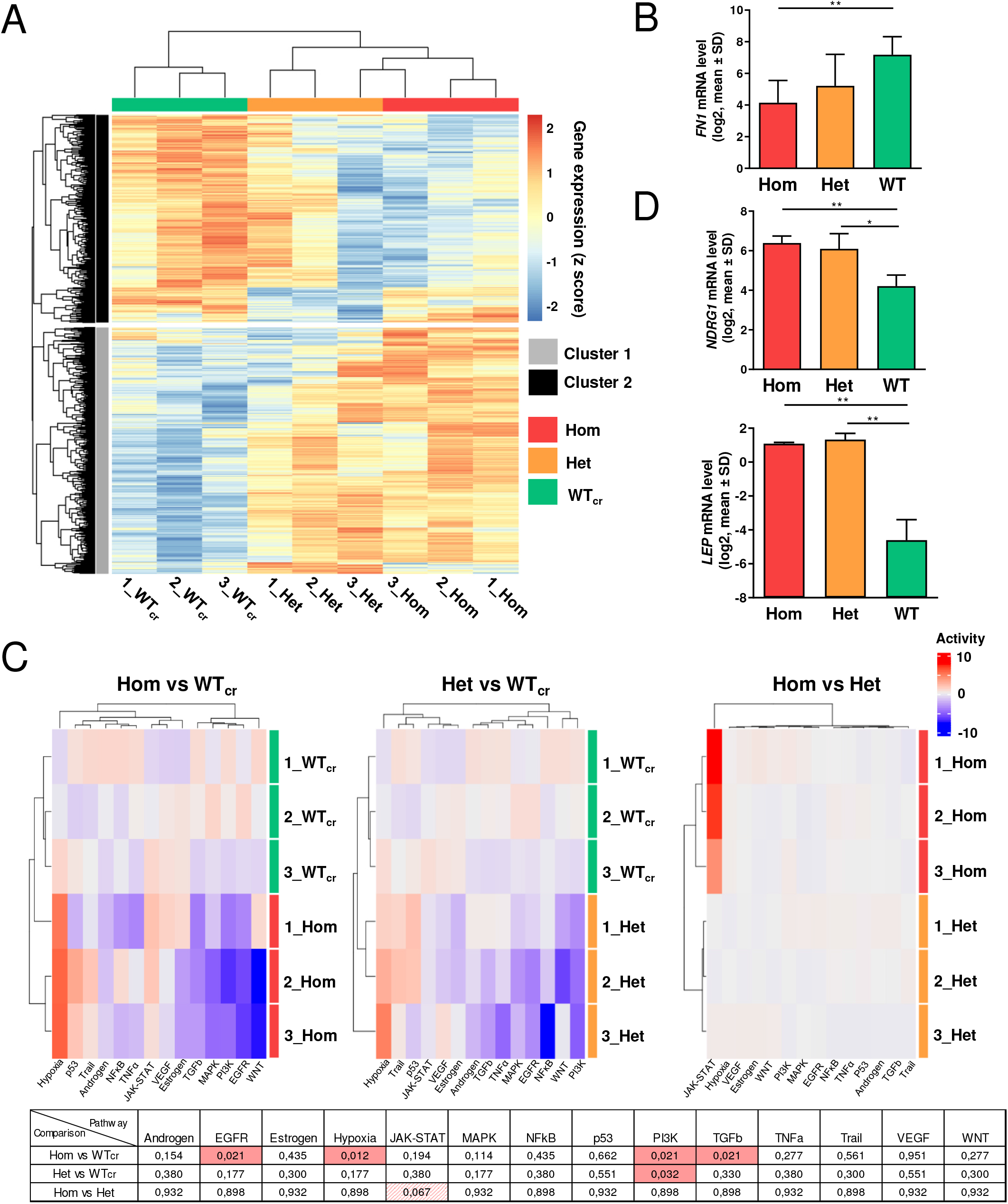
RNAseq experiments with iPS cell-derived MKs of *CALR*ins5-mutated clones. **(A)** Heat map for unsupervised hierarchical clustering of RNAseq data of iPS cell-derived MKs with homozygous (Hom) or heterozygous (Het) *CALR*ins5 mutation or unmutated (WT_cr_) MKs. Numbers represent independent experiments. Unsupervised clustering was performed. Cluster 1 and cluster 2 are indicated in grey and black, respectively. **(B)** Gene expression profile of *FN1* in iPS cell-derives MKs of RNAseq analysis. Analysis was performed for three independent experiments. \*\**p*<0.01. **(C)** Pathway Responsive genes (PROGENy) of multiple comparisons of Hom, Het and WT_cr_ iPS cell-derived MKs. Upregulated and downregulated pathways were shown in red and blue, respectively. Adjusted p-values from multiple comparisons are shown in the table. Significant values (p.adjusted<0.05) are highlighted in red. **(D)** Gene expression profile of *NDRG1* and *LEP* in iPS cell-derives MKs of RNAseq analysis. Analysis was performed for three independent experiments. \**p*<0.05, \*\**p*<0.01.

## Discussion

In the present study, we generated patient-specific iPS cell lines harboring *CALR*del52, ins5 or del31 mutations, and the iPS cell-based model was applied to study the impact of *CALR* mutant zygosity on hematopoietic and more precisely on megakaryocytic differentiation at the clonal level. Patient-derived iPS cells carrying heterozygous *CALR* mutations have been recently described, but detailed comparisons of type 1 and type 2 *CALR* mutations with focus on their zygosity and detailed characterization of derived disease-driving MKs are missing (Gomez Limia et al., 2018, 2017; Secardin et al., 2021; Takei et al., 2018). To obtain isogenic *CALR*-repaired clones for *CALR*ins5 and del52-mutated iPS cells, CRISPR/Cas9 technology was used to correct the mutations. Until now, the repair of *CALR*ins5 mutations was reported at a conference abstract (Wang et al., 2018), and a CRISPR/Cas9 approach was used to model *CALR* mutations in murine cell lines (Abdelfattah and Mullally, 2018). To our knowledge, the repair of *CALR*del52 mutations in MPN-specific iPS cells was not reported, yet. The absence of further MPN-related mutations of clinical significance was confirmed by NGS in our *CALR*-mutated iPS cell clones. Hence, our findings are solely attributed to the *CALR* mutations.

MPO deficiency was demonstrated in homozygous *CALR*ins5 iPS cell-derived CD15+ cells and, for the first time, also in homozygous *CALR*del52 cells. Loss of MPO activity in MPN patients carrying homozygous CALR mutations was described by Theocharides and colleagues (Theocharides et al., 2016), and we were able to restore MPO activity by CRISPR/Cas9 gene repair, demonstrating the recovery of the chaperon function of CALR. The rescue of CALR WT characteristics was underlined by the fact that WT_cr_ and HD control iPS cells generated equal levels of MKs. Importantly, the iPS cell-derived MKs exhibited structural features of primary MKs such as formation of DMS, granules, multi-lobulated nuclei and proplatelets.

Megakaryocytic hyperplasia was reported for all types of MPNs and represents a diagnostic criterion of the World Health Organization (Barbui et al., 2018; Swerdlow et al., 2008; Zingariello et al., 2020). Our data demonstrate that both *CALR*ins5- and del52-mutated iPS cells gave rise to more MKs than WT_cr_ and HD control iPS cells. In particular, homozygous mutant clones produced higher numbers of MKs followed by heterozygous clones. Thus, the mutated CALR phenotype demonstrated in our study recapitulates the phenotype found in MPN patients and the data of Kollmann and colleagues demonstrating that the ectopic expression of CALRdel52 or CALRins5 mutants in human CD34+ HSPCs lead to enhanced megakaryopoiesis and platelet production (Kollmann et al., 2017). Furthermore, knock-in mice expressing homozygous *CALR*ins5 or del52 mutations exhibited stronger ET- and PMF-related phenotypes over heterozygous CALR-mutant mice, demonstrating that zygosity of the *CALR* mutation has an impact on disease development (Benlabiod et al., 2020). Araki and colleagues described mutant CALR protein homomultimers that are presumed to bind and activate the TPO receptor (Araki et al., 2018). This observation is confirmed by our data as a strong bias towards the MK lineage and enhanced MK maturation was observed for homozygous *CALR*-mutated iPS cells. Hence, we assume that mutant CALR homomultimers are formed more excessively in homozygous clones than in heterozygous clones.

Importantly, we demonstrated in our human iPS cell-based model that the *CALR* mutation led to the amplification of the MK lineage in an TPO-independent manner while WT and HD control cells showed pronounced MK population only in the presence of TPO. Of note, additional treatment of TPO did not further increase MK generation of *CALR*-mutated cells. This might be due to the reported oncogenic interaction of the TPO receptor and mutant CALR before reaching the cell surface (Masubuchi et al., 2020; Pecquet et al., 2019), already sufficiently occupying the TPO binding site. However, Secardin and colleagues observed an increase of CFU-MKs, generated from iPS-cell-derived CALR-mutated CD34+ CD43+ progenitors, in the presence of TPO (Secardin et al., 2021). These differences may be clone-dependent or exemplifies differences between liquid cultures and collagen/fibrin-based medium.

The observed amplification of the MK lineage in *CALR*-mutated cells raises the question whether it relies on enhanced proliferation or accelerated maturation of the HSPCs or MKs. Immunohistochemical staining of MKs showed a higher expression of proliferative *Ki67* compared to controls (Malherbe et al., 2016). However, stronger *Ki67* gene expression was not confirmed in our iPS cell-derived *CALR*-mutated MKs, but in the *CALR*ins5 CD34+ population. Here, we observed a higher number of colonies in the CFU assays for *CALR*ins5-mutated cells and higher cell numbers on day 14 of differentiation in the *CALR*-mutated background compared to unmutated cells. Those data demonstrate that *CALR* mutations (especially *CALR*ins5) impact on proliferation capacity of immature hematopoietic cells and MK differentiation.

While *CALR* mutations did not impact on MK proliferation, we showed that homozygous and heterozygous *CALR* mutations induced an accelerated maturation of MKs. Surface expression of CD42b, which is a marker for megakaryocytic maturation (Sim et al., 2017), was elevated on mutated MKs in comparison to WT_cr_ and HD control MKs at an early time point of differentiation. Furthermore, higher expression of *NFE2* in homozygous and heterozygous *CALR*ins5 MKs compared to WT_cr_ MKs was observed. Moreover, we found that *CALR*ins5 homozygous mutated MKs exhibited a higher level of granularity compared to cells harboring unmutated or heterozygous CALRins5. These findings are supported by Sim et al, that identified different maturation stages in MKs corresponding to their granularity (Sim et al., 2017). Taken together, our data demonstrate that *CALR* mutations cause an accelerated megakaryopoiesis in iPS cell-derived MKs rather than affecting megakaryocytic proliferation. In future work, our model could be used to study the relationship of accelerated and enhanced megakaryopoiesis and platelet production/release in *CALR*-mutated MKs.

We confirmed the upregulation of typical megakaryocytic genes such as *NFE2, FLI1* and *VWF* in our iPS cell-derived MKs by RT-qPCR analysis and proceeded to an in-depth analysis of the global gene expression profile by RNAseq. Recent studies reported on MPN-related transcriptome analysis of *in vitro* differentiated MKs from primary *CALR*-mutated CD34+ (El-Khoury et al., 2020) or single-cell RNAseq of primary CD34+ cells from MF patients with either *JAK2*V617F or *CALR* mutation (Psaila et al., 2020). In contrast to the first mentioned study, we were able to relate differences of the megakaryocytic transcriptome to the underlying *CALR* genotype by analyzing a clonal cell population. Therefore, our model with defined mutational background more precisely elucidates mechanisms of aberrant megakaryopoiesis in MPN.

Our RNAseq data identified elevated interferon signaling and leptin gene expression in *CALR*-mutated MKs. Leptin functions as a hormone and adipokine predominantly expressed by adipocytes (Cava and Matarese, 2004). However, a role of leptin in proinflammatory processes was reported, as it is able to activate human peripheral blood B cells to induce the secretion of cytokines such as TNF-α, IL-6 and IL-10 via JAK2/STAT3 and p38MAPK/ERK1/2 signaling pathways (Agrawal et al., 2011). In liver fibrosis, leptin promotes the proliferation and secretion of ECM molecules in myofibroblasts (activated hepatic stellate cells) via the leptin receptor (Vivoli et al., 2016). Also, in BM fibrosis, leptin receptor-positive mesenchymal stromal cells were identified to be responsible for collagen fiber generation and deposition (Decker et al., 2017). Hence, it is conceivable that megakaryocyte-produced leptin may be implicated in myofibroblast formation in MPN and contribute to the inflammatory environment in the BM niche.

Of note, the expression of ECM proteins was downregulated in *CALR*-mutated MKs, compared to control. The composition of ECM in the BM is essential for megakaryocytic differentiation and maturation (Noetzli et al., 2019). Synthesis of ECM components in MKs was described to be regulated via autocrine TGF-β1 signaling (Abbonante et al., 2016). Our PROGENy analysis revealed that TGF-β signaling was significantly downregulated in homozygous *CALR*-mutant MKs, which may explain the downregulation of ECM genes in *CALR*-mutated MKs. However, in the above-mentioned study (Abbonante et al., 2016), *in vitro* experiments of CD34+-derived MKs from *JAK2*V617F-mutated PMF patients were performed, showing a stronger activation of TGF-β signaling and, thus, increased expression of ECM components such as fibronectin and type IV collagen compared to control. However, the allele burden of the *JAK2*V617F mutation in the bulk of primary CD34+ cells was not specified. In contrast, ECM gene expression in our clonal iPS cell-derived MKs was solely correlated to the *CALR* mutation.

In addition, the hypoxia signaling pathway was strongly upregulated in *CALR*-mutated vs unmutated MKs. In solid tumors, it is well established that hypoxia represents a key feature for tumor survival and development (Thomlinson and Gray, 1955), and the BM stem cell niche represents an environment of relatively low oxygen level (Irigoyen et al., 2017). The major regulator of cellular response to hypoxia is the hypoxia-inducible factor 1 (HIF-1a) (Semenza, 2009). Importantly, HIF-1a promotes survival of *JAK2*V617F-positive cells and was discovered to be a promising therapeutic target (Baumeister et al., 2020). Moreover, elevated protein levels of HIF-1a were recently described in blast phase MPN samples, suggesting a role of HIF-1a in MPN disease progression (Marinaccio et al., 2021). In addition, El-Khoury et al identified hypoxia as a deregulated pathway in primary CD34+ cells differentiated towards MKs from MPN-patients (El-Khoury et al., 2020). Hence, our data support the notion that hypoxia-related factors may be promising targets in *CALR*-mutated MPNs.

Collectively, our study describes a comprehensive iPS cell system for evaluating mutant *CALR*-induced phenotypes in MPN with a clonal genetic background, which is an valuable new tool for clinical drug screening and development of personalized therapies. We identified accelerated megakaryopoiesis and the upregulation of leptin expression as well as hypoxia and interferon signaling pathways as potential pathogenic mechanisms in *CALR*-mutated MKs.

## List of abbreviations

CALR: calreticulin
CFU: colony forming unit
DEGs: differentially expressed genes
DMS: demarcation membrane system
EB: embryoid body
ET: essential thrombocythemia
FCS: forward scatter
iPS: induced pluripotent stem
MF: myelofibrosis
MKs: megakaryocytes
MPL: thrombopoietin receptor
MPN: myeloproliferative neoplasm
MPO: myeloperoxidase
OCS: open canalicular system
PBMCs: peripheral blood mononuclear cells
PMF: primary myelofibrosis
PV: polycythemia vera
SSC: side scatter
TEM: transmission electron microscopy
TPO: thrombopoietin
HSPCs: hematopoietic stem and progenitor cells

## Declarations

### Ethics approval and consent to participate

Peripheral blood mononuclear cells (PBMCs) were obtained from MPN patients carrying *CALRdel52* or *CALRins5* mutation at the Department of Hematology at Zurich University, and from a *CALR* del31 mutated MPN patient at the centralized Biomaterial Bank in Hospital RWTH Aachen, University. Healthy donor PBMCs were provided from the Transfusion Medicine in the University Hospital RWTH Aachen. All PBMCs were donated after written informed consent, as approved by the ethics committees in Zurich and the Medical Faculty of RWTH Aachen (EK127/12, EK 206/09, and EK099/14).

### Consent for publication

All authors have signed the form of consent to publication.

### Competing interests

SK reports funding from Novartis, Bristol-Myers Squibb, Janssen/Geron; advisory board honoraria from Pfizer, Incyte, Ariad, Novartis, AOP Pharma, BMS, Celgene, Geron, Janssen, CTI, Roche, Baxalta, Sanofi; patent for BET inhibitor at RWTH Aachen University; honoraria from Novartis, BMS, Celgene, Geron, Janssen, Pfizer, Incyte, Ariad, Shire, Roche, AOP Pharma; and other financial support (e.g. travel support) from Alexion, Novartis, BMS, Incyte, Ariad, AOP Pharma, Baxalta, CTI, Pfizer, Sanofi, Celgene, Shire, Janssen, Geron, Abbvie, Karthos.

## Material and methods

### Generation of MPN patient-specific and healthy donor iPS cells

Peripheral blood mononuclear cells (PBMCs) were obtained from MPN patients carrying a *CALRdel52* or *CALRins5* mutation at the Department of Hematology at Zurich University, and from a *CALR*del31 mutated MPN patient at the centralized Biomaterial Bank in Hospital RWTH Aachen, University. HD PBMCs were provided from the Transfusion Medicine in the University Hospital RWTH Aachen. All PBMCs were donated after written informed consent, as approved by the ethics committees in Zurich and the Medical Faculty of RWTH Aachen. PBMCs were reprogrammed using a CytoTune iPS 2.0 Sendai Reprogramming Kit (Thermo Fisher Scientific, Waltham, MA; USA) according to the manufacturer’s instructions. After reprogramming, individual iPS cell colonies were picked and screened for *CALR* genotypes by PCR or allele-specific PCR.

### Next generation sequencing

For the detection of mutations from gDNA by next generation sequencing (NGS), two panels individually designed and validated for routine hematology diagnostic were used. Either 250ng or 75ng of gDNA were used for library preparation with either a Truseq Custom Amplicon Kit (Illumina, San Diego, USA) or an Ampliseq for Illumina Custom Panel (Illumina) covering the relevant regions of either 31 (*ABL1, ASXL1, BARD1, CALR, CBL, CHEK2, CSF3R, DNMT3A, ETNK1, ETV6, EZH2, IDH1, IDH2, JAK2, KIT, KRAS, MPL, NFE2, NRAS, PDGFRA, PTPN11, RUNX1, SETBP1, SF3A1, SF3B1, SH2B3 (LNK), SRSF2, TCF12, TET2, TP53, U2AF1*)(Kirschner et al., 2018) or 32 genes (*ABL1, ASXL1, BRAF, BTK, CALR, CBL, CSF3R, CXCR4, DNMT3A, ETNK1, EZH2, FLT3, IDH1, IDH2, JAK2, KIT, KRAS, MPL, MYD88, NF-E2, NPM1, NRAS, PTPN11, RUNX1, SETBP1, SF3B1, SH2B3, STAT5B, TET2, TP53, U2AF1, WT1*) associated with hematologic malignancies. The final libraries were sequenced with 2×250bp on a MiSeq (Illumina). The MiSeq onboard software was used (Real time analysis software v1.18.54, Illumina) for demultiplexing and FastQ file generation. Alignment and variant calling were performed with the SeqNext-Module of the SeqPilot-Software (JSI medical systems, Version 4.4.0 Build 509). Variants were called with a bidirectional frequency of >5% (JAK2 V617F and KIT D816V >1%) and reviewed manually.

### Cytogenetic analysis

Patient-derived iPS cell clones were seeded on matrigel coated 25 cm^2^ cell culture flasks. Normal chromosomal constitution was verified by conventional karyotyping of iPS cell clones by means of GTG banding at 400 to 550 band level. Metaphase spreads were prepared using standard procedures of blocking cell division at metaphase, hypotonic treatment, and methanol/acetic acid fixation (3:1). The banding techniques included the use of a trypsin pretreatment (GTG-banding) carried out according to standard protocols. Microscopy was performed with Axioplan fluorescence microscope (Carl Zeiss) and IKARUSTM digital imaging systems (MetaSystems, Altlussheim, Germany). An average of 20 mitoses were analyzed for each clone.

### Immunofluorescence staining

iPS cell colonies were cultured on Matrigel coated coverslips in 4-well-plates. Cells were fixed with 4 % paraformaldehyde (Sigma Aldrich) and blocked with goat serum (Merck Millipore, Darmstadt, Germany). In order to detect the expression of pluripotent markers and mutant CALR, cells were incubated with following primary antibodies at 4 °C overnight: TRA-1-60 (Merck Millipore), OCT3/4 (H-134, Santa Cruz, CA, USA) or mouse monoclonal antibody CAL2 (Dianova, Hamburg, Germany). Cells were washed three times with PBS on the next day and incubated with corresponding secondary antibodies goat anti-rabbit IgG (H+L) FITC (Thermo Fisher Scientific), goat anti-mouse IgM (H) Alexa Fluor 594 (Thermo Fisher Scientific) or goat anti-mouse IgG Alexa Fluor 647 (for CALR mutant, clone poly4053, BioLegend, San Diego, CA, USA) for 1 h in the dark. Nuclei were stained with DAPI (Vector Laboratories, Burlingame, CA, USA) and coverslips were mounted with Dako Fluorescence Mounting Medium (Dako, Jena, Germany). Fluorescent image analysis was performed using an Axiovert 200 microscope (Carl Zeiss, Jena, Germany) and IPLab Spectrum software (BD, Franklin Lakes, USA).

In order to detect localization of WT and mutand CALR in MKs, cells were fixed with 3,7 % PFA, washed with PBS and permeabilized with 0,1 % Trition-X and 2,5 % BSA. Afterwards cells were incubated with detection reagent for ER-staining and Hoechst (Enzo Life Sciences, Farmingdale,USA) for 20 min at 37°C. Cells were washed with PBS and blocked wit 5 % BSA in PBS. Cells were incubated with the following primary antibodies at 4 °C overnight: polyclonal antibody CD42b (Thermo Fisher Scientific) and mutant CALR antibody (Dianova) or WT CALR monoclonal antibody (Novus, Littleton, USA). On the next day, cells were washed with PBS and incubated with corresponding secondary antibodies goat anti-rabbit IgG Alexa Fluor 555 (H+L) and goat anti-mouse IgG (H+L) Alexa Fluor 647(all Thermo Fisher Scientific) for 1 h at RT. Coverslips were mounted with Fluorescence Mounting Medium (Dako, Carpinteria, USA). Fluorescent image analysis was performed using a confocal Scanning Microscope (LSM 710) microscope and Zeiss 2012 software (both Carl Zeiss, Jena, Germany).

### RNA isolation and qPCR

RNA of iPS cells and iPS-cell derived CD34+ HSCS and CD61+ MKs was isolated using the RNeasy Mini Kit (QIGAEN GmbH, Hilden, Germany) according to manufacturer’s protocol. RNA (500 ng) was used for cDNA synthesis. Quantitative RT-PCR was performed using the 7500 Fast Real-time PCR System (Applied Biosystems by Life technologies, Paisley, UK) with the SYBR Select Master Mix for CFX (Applied Biosystems). The sequences of primers used for-qPCR are listed in Table S7. All primers were purchased from Eurofins-MWG biotech (Ebersberg, Germany). The mRNA expression level of the target gene is determined in % of *GAPDH* or *MT-ATP6*.

### SDS-Page and Western blot

SDS-Page and Western blot analysis were conducted as previously described (Han et al., 2016). Primary and secondary antibodies are listed in Table S6.

### CRISPR/Cas9n-mediated repair of homozygous *CALRins5* and *CALRdel52* mutation in iPS cells

The Alt-R™ CRISPR-Cas system (Integrated DNA Technologies IDT, Coralville, USA) was used to efficiently correct homozygous *CALRins5* or *CALRdel52* mutations in MPN patient-specific iPS cells. For precise editing the Alt-R® S.p HiFi Cas9 Nuclease V3 was used together with a single-stranded donor template to repair the double strand break by the cellular repair mechanisms of homology directed repair (HDR). In brief, guide RNA composing of crRNA and tracrRNA was combined with HiFi. Cas9 Nuclease to assembly the CRISPR-Cas9 ribonucleoprotein complex. Singularized iPS cells were gently mixed with the ribonucleoprotein complex, single-stranded donor template, and electroporation enhancer. Nucleofection was performed using the 4D NucleofectorTM X-Unit and the P3 Primary Cell 4D-NucleofectorTM X, Kit S (both from Lonza, Basel, Switzerland). HDR was stimulated by HDR enhancer (IDT, Coralville, USA). After nucleofection, iPS cells were seeded on previously coated Laminin 521 (Biolamina, Sundbyberg, Sweden) plates to support cell viability in StemMACS™ iPS-Brew XF (Miltenyi Biotec, Bergisch Gladbach, Germany) supplemented with 10 µM of Rock inhibitor Y-27632 and 1x CloneR (Stemcell Technologies). Repair of *CALR*ins5 and *CALR*del52 mutation was verified by allele-specific or flanking PCR, respectively, and proofed by Sanger sequencing and Western blot. Possible off-target effects were excluded by PCR reaction of off-target sites and subsequent Sanger sequencing. Sequences of crRNA, donor template and primers are provided in Table S2 and 3.

### Differentiation of iPS cells into hematopoietic progenitors and myeloid subsets with the “EB-based” protocol

iPS cells were cultured on irradiated mouse embryonic fibroblast feeder layer in KnockOut-DMEM based iPS cell medium. More detailed information regarding cell culture conditions can be found in Table S8.

For the generation of iPS cell-derived CD34+ HSCs, CALR iPS cells were subjected to mesoderm commitment and hematopoietic differentiation with an embryonic body (EB)-based protocol modified from the differentiation method described by Kovarova and colleagues (Kovarova and Koller, 2012). CD34+ HSCs were cultured and differentiated into myeloid subsets in the StemPro 34 SFM medium (Thermo Fisher Scientific) based progenitor medium at the maximum density of 1×10^6^ cells/ml. Details of EB and HSCs differentiation media are provided in Table S8.

### Cell morphology analysis

Phase contrast images of iPS cell generation and differentiation were obtained with EVOS FL microscope (Thermo Fisher Scientific). To characterize the morphologies of iPS cell-derived precursors and myeloid subsets, cells were spun onto slides with Shandon Cytospin 4 cytocentrifuge (Thermo Fisher Scientific) and stained with Diff Quik solution 1 and 2 (Medion Diagnostics, Düdingen, Switzerland) after methanol fixation. Images were acquired using a Leica DMRX microscope and Leica Application Suite software (Leica Microsystems).

### MPO cytochemical staining

In order to evaluate MPO functional activity, iPS cell-derived hematopoietic cells at differentiation day 15 were centrifuged onto slides and fixed with LEUCOGNOST® Fixing Mixture. The fixed cells were stained with LEUCOGNOST® POX Kit according to the manufacturer’s instructions. Counterstaining of nucleus was performed using Mayer’s hemalum solution and slides were mounted with Kaiser’s glycerol gelatin (the Kit and all reagents were obtained from Merck Millipore). Images were acquired as described above.

### Differentiation of iPS cells into hematopoietic stem cells and myeloid subsets with the “spin-EB” protocol

Feeder-free iPS cell culture was maintained on matrigel coated 6-well plates and routinely passaged with 1 ml accutase or 0.5 mM EDTA (Thermo Fisher Scientific). To enhance single cell survival, 10 µM ROCK inhibitor Y-27632 was added to the maintenance culture medium StemMACs iPS Brew XF for 24 h after seeding. iPS cell medium was changed daily.

Human iPS cells were differentiated into HSPCs, MKs and erythrocytic cells adapted from the differentiation protocol by Liu et. al (Liu et al., 2015). Briefly, iPS cells were seeded into U-bottom-shaped 96-well plates with a density of 3,000 cells/well in cytokine supplemented serum-free medium (SFM). To allow spheroid formation, plates were spun at 380 g for 5 min. From day 2 to day 8, cells were cultured in SFM with 10 ng/ml VEGF (Miltenyi Biotec), 10 ng/ml BMP4 (Miltenyi Biotec), 10 ng/ml bFGF (Peprotech, Hamburg, Germany) and SCF (0.5 % supernatant of SCF secreting CHO KLS cells). From day 8 onwards, BMP4 and VEGF were removed from the medium. On day 11, 20 ng/ml TPO (Miltenyi Biotec) was added to the medium. On day 14, cells were harvested and filtered through a 100 µm filter to separate the EBs from the suspension cells. Single cells were analyzed by flow cytometry and purified for CD61+ and CD34+ cell fractions. More detailed information on SFM Medium is given in Table S8.

### Flow cytometry analysis

To evaluate cell surface progenitor and lineage specific markers, single cells were analyzed on day 10, day 12 and day 14 of “spin-EB” differentiation and on day 10 and day 15 of “EB-based” differentiation. Cells were harvested and passed through a 100 μm cell strainer, centrifuged at 350 g for 5 min. After washing with ice cold FACS buffer, single cells were incubated with lineage-specific antibodies at 4°C for 30 min. Stained cells were resuspended in 300 μl FACS buffer and analyzed on a FACS Canto II (BD). All staining and washing steps were performed in FACS buffer consistent of PBS supplemented with 2 mM EDTA and 2 % BSA (PAN Biotech, Aidenbach, Germany).

To identify the intracellular MPO expression, differentiated hematopoietic cells at day 15 of “EB-based” differentiation were fixed and permeabilized using Fix & Perm Cell Permeabilization Kit (Thermo Fisher Scientific). Cells were firstly stained for cell surface markers CD15 and CD45 for 20 min at room temperature (RT), followed by intracellular MPO staining for 20 min at RT after fixation and permeabilization. Stained cells were evaluated by Gallios flow cytometer (Beckman Coulter, Krefeld, Germany). All data were analyzed by FlowJo™ (version 10, Oregon, USA).

Full list of antibodies is summarized in Table S5. In each experiment matched Ig isotype controls were used to set background fluorescence.

### Purification of CD34+ HSPCs and CD61+ megakaryocytes by magnetic activated cell sorting

On day 25-35 of “EB-based” differentiation and on day 14 of “spin-EB” differentiation anti-CD34 and anti-CD61 MicroBeads (Miltenyi Biotec, Bergisch Gladbach, Germany) were used to separate HSPCs and MKS, respectively. Purification of cells was performed according to the protocol provided by the manufacturer. In brief, single cell suspension was incubated with CD61 or CD34 MicroBeads for 15 or 30 min, respectively, to label the cells magnetically. Cell suspension was loaded onto a LS column installed in a magnetic field. After washing three times with MACS buffer consisting of PBS supplemented with 5 % FCS (Thermo Fisher Scientic) and 2 mM EDTA to remove unbounded cells, the column was removed from the magnetic field and positively selected cell fraction was eluted with 5 ml of MACS buffer from the column. After centrifugation at 300 g for 10 min, purified cells were counted and further processed. CD61+ MKs and CD34+ HSPCs were either snap frozen for RT-qPCR or used for transmission electron microscopcy or seeded in Colony-Forming Unit Assay, respectively.

### Colony-forming unit assay of CD34+ HSPCs

Purified CD34+ cells were seeded in MethoCult™ (Stemcell Technologies) supplemented with 20 ml IMDM (Thermo Fisher Scientific), 50 ng/ml hSCF, 10 ng/ml hIL-3. 10 ng/ml hGMSCF and 14 ng/ml hEPO (all ImmunoTools Friesoythe; Germany) in a cell density of 5,000 cells/ml. After 10-12 days of culture at 37° C, colonies were identified and counted based on morphology using light microscopy (Motic, Barcelona, Spain).

### Transmission electron microscopy

iPS cell-derives CD61+ megakaryocytes at day 14 of “spin-EB” differentiation were prepared as described previously. Purified MKs were washed with PBS and fixed with 3 % glutaraldehyde for at least 2 h at RT, followed by embedding in 5 % low melting agarose (Merck). Gelatinated blocks were washed in 0.1 M Soerensen’s phosphate buffer (Merck) and post-fixed in 17 % sucrose buffer (Merck) containing 1 % OsO4 (Roth, Karlsruhe, Germany). Subsequently, specimens were dehydrated performing an ascending ethanol series repeating the last step for three times (30, 50, 70, 90, and 100 % ethanol; 10 min each step). Dehydrated samples were consecutively incubated in propylene oxide (Serva, Heidelberg Germany) for 30 min, in a mixture of EPON resin (Serva) and propylene oxide (1:1) for 1 h, and in pure EPON for 1 h. EPON polymerization was conducted at 90 °C for 2 h. Ultrathin sections of 70−100 nm were cut with a Reichert Ultracut S ultramicrotome (Leica, Wetzlar, Germany) equipped with a diamond knife (Leica) and picked up on copper−rhodium grids (Plano, Wetzlar, Germany). Contrast was enhanced by staining with 0.5 % uranyl acetate and 1 % lead citrate (both EMS, Munich, Germany). Samples were viewed at an acceleration voltage of 60 kV using a Zeiss Leo 906 (Carl Zeiss, Jena, Germany) transmission electron microscope. Image processing and analysis was performed using ImageJ (Schneider et al., 2012).

### Preparation of samples for RNA Sequencing

RNA of purified CD61+ MKs was isolated as described in Supplementary Information. Quality was verified by TapeStation 4200 (Agilent, Santa Clara, USA) to determine RNA integrity number (RIN). RNA concentration was measured with Fluorometer Quantus™ (Promega, Fitchburg, USA). According to manufacturer’s protocol, ribosomal RNA was depleted using NeBNext® rRNA Depletion Kit, prior to library preparation with NeBNext®Ultra™II Directional RNA Library Prep Kit for Illumina (both New England BioLabs, Ipswich, USA) using an RNA input of 100 ng per sample (New England BioLabs, Ipswich, USA). Subsequently, samples were sequenced in paired end reads (2×76 bp, dual indexed) on two NextSeq High Output Kits v2.5 (150 cycles) on a NextSeq 500 instrument (both Illumina, San Diego, USA) to have sufficient reads for data analysis.

### Statistical analysis

Graphical display and statistical analysis were performed with Prism 7 (GraphPad, San Diego, California, USA). Unless otherwise stated, all experiments were performed in triplicates. For each *CALR* genotype, two different clones were used, except for heterozygous *CALRdel52* clone, where only one clone was generated after reprogramming. Comparisons among two groups were performed by using unpaired Student *t* test. For multiple group comparisons, *ANOVA* with Bonferroni posttest was performed. *p* values <0.05 were considered statistically significant (\**p*<0.05, \*\**p*<0.01, \*\*\**p*<0.001).

### Online supplemental material

Fig S1 A shows karyotype analysis of patient-derived iPS cell cones. Fig S1, B and C display the alignment of iPS cell clones after CRISPR/Cas9 repair of *CALR*-mutated maternal clone with CALR WT sequence. Fig. S1, D and E show sequencing analysis of potential CRISPR/Cas9 off-target products. Fig. S2, A and B show gating strategy of flow cytometry analysis exemplarily. Fig. S2 C shows number of colonies counted in a CFU assay, related to Fig. 3 D. Fig. S2 D shows *Ki67* expression of CD34+ HSPCs. Fig. S2 E shows the quantification of erythrocytic cells analyzed by flow cytometry. Fig. S2 F demonstrates *Ki67* expression of CD61+ MKs. Fig. S3, A and B show immunofluorescence analysis of mutant and WT CALR in iPS-cell derived MKs. Fig. S4 shows gene ontology (GO) analysis of clusters identified in a heat map of DEGs shown in Fig. 7 A. Fig. S5, A, B and C show volcano plots of DEGs compared among *CALR* genotypes. Fig. S7 D shows validation of genes identified in Fig. 7 D by RT-qPCR.

Table S1 summarizes details of reprogrammed patients’ samples and generated iPS cell clones. Table S2 lists guide RNA used for CRISPR/Cas9 mediated *CALR* repair. Table S3 shows donor templates used for homology directed repair. Table S4 displays MPN-related mutations and polymorphisms of *CALR* iPS cell clones. Table S5 lists antibodies used for flow cytometry. Table S6 lists primary and secondary antibodies used for Western blot and Immunofluorescence staining. Table S7 summarizes Primers used for genotyping and RT-qPCR. Table S8 summarizes media compositions.

## Acknowledgements

We thank Reinhild Herwartz, Melanie Baumann and Lucia Vankann for technical assistance of MPO cytochemical staining and flow cytometry analysis, Ulla Gollan for cytogenetic analysis. We thank Sabrina Ernst, Carmen Schalla, and Prof. Dr. rer. nat. Gerhard Müller-Newen for technical support on immunofluorescence staining experiments. Biomaterial samples were provided by the RWTH centralized Biomaterial Bank Aachen (RWTH cBMB, Aachen, Germany) in accordance with the regulations of the biomaterial bank and the approval of the ethics committee of the medical faculty, RWTH Aachen. This work was in part supported by the Flow Cytometry Facility, a core facility of the Interdisciplinary Center for Clinical Research (IZKF) Aachen within the Faculty of Medicine at RWTH Aachen University. This work was supported by the Confocal Microscopy Facility, a Core Facility of the Interdisciplinary Center for Clinical Research (IZKF) Aachen within the Faculty of Medicine at RWTH Aachen University. The study was supported by the Genomics Facility, supported by a grant from the Interdisciplinary Center for Clinical Research within the Faculty of Medicine at RWTH Aachen University.

## Funding

This work was supported in part by funds from the German Research Foundation (Deutsche Forschungsgemeinschaft DFG CH1509/1-1, KO2155/7-1, ZE432/10-1 and GE 2811/4-1) as part of the clinical Research Unit CRU344, a research grant from the Deutsche José Carreras Leukämie-Stiftung (DJCLS 16R/2017) to SK, and a research grant from the German Research Foundation (DFG KO2155/6-1) to SK. LH was supported by the National Natural Science Foundation of China (No. 82000133). MAST was financed by CAPES-Alexander von Humboldt postdoctoral fellowship (99999.001703/2014-05). AT was supported by the Prof. Dr. Max Cloëtta foundation. Parts of this work were generated within the PhD thesis project of KO and LH.

## Author contributions

Conceptualization: K. Olschok, L. Han, M.A .S. Toledo, N. Chatain, and S. Koschmieder;

Data curation: M. Graßhoff, I.G. Costa, A. Maurer, and J. Baumeister;

Formal analysis: K. Olschok, L. Han, M. Graßhoff, I.G. Costa, A. Maurer, and J. Baumeister;

Funding acquisition: N. Chatain, and S. Koschmieder;

Investigation: K. Olschok, L. Han, M. A .S. Toledo, J. Böhnke, A. Maurer, H. M. Schüler, E. M. Buhl, K. Pannen, S. Gupta, and P. Boor;

Methodology: K. Olschok, L. Han, M. A. S. Toledo, and J. Böhnke

Project administration: K. Olschok, L. Han, M.A .S. Toledo, N. Chatain, and S. Koschmieder;

Resources: A. Theocharides, M. Kalmer, D. Gezer, and M. Zenke;

Software: M. Graßhoff, and I. G. Costa;

Supervision: M .A .S. Toledo, N. Chatain, and S. Koschmieder;

Validation: K. Olschok, and L. Han;

Visualization: K. Olschok, L. Han, M. A. S. Toledo, and M. Graßhoff

Writing – original draft: K. Olschok, and L. Han;

Writing – review & editing: K. Olschok, L. Han, M. A .S. Toledo, J. Böhnke, M. Graßhoff, I.G. Costa, A. Theocharides, A. Maurer, H. M. Schüler, E. M. Buhl, K. Pannen, J. Baumeister, M. Kalmer, S. Gupta, P. Boor, D. Gezer, T. H. Brümmendorf, N. Chatain, and S. Koschmieder;

All authors approved the final version of the manuscript.

Kathrin Olschok and Lijuan Han have contributed equally to this work.

Nicolas Chatain and Steffen Koschmieder share last authorship.

## Corresponding authors

Correspondence to Nicolas Chatain and Steffen Koschmieder.

