## Supplementary data for "CALR frameshift mutations in MPN patient-derived iPS cells accelerate maturation of megakaryocytes"

[#]Shared first authorship

*Shared senior authorship

**Correspondence:** Prof. Dr. med. Steffen Koschmieder, Department of Hematology, Oncology, Hemostaseology, and Stem Cell Transplantation, Faculty of Medicine, RWTH Aachen University, Pauwelsstr. 30, D-52074 Aachen, Germany, Phone: +49-241-8036102;

Supplemental Figures

Figure S1:

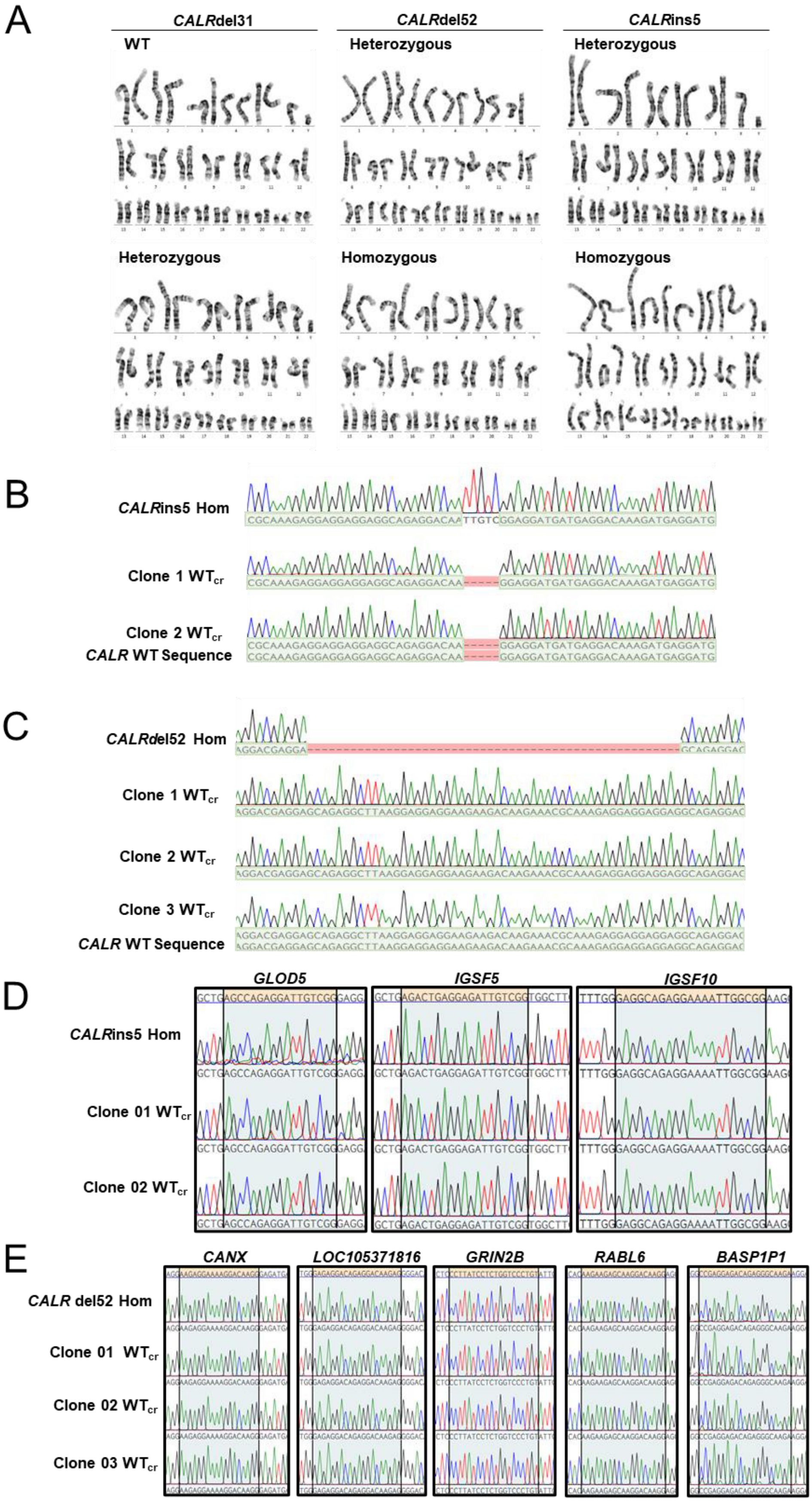

**Figure S1. Proof of CRISPR/Cas9 engineered *CALR* WT clones.**

**A.** Karyotype analysis of patient-derived iPS cell clones. **B.** Sequence alignment of maternal homozygous *CALR*ins5 clone with resulting $WT_{cr}$ clones 01 and 02 to reference CALR WT sequence. Successful alignment and mismatch are shown in green and red, respectively. **C.** Sequence alignment of maternal homozygous *CALR*del52 clone with resulting $WT_{cr}$ clones 01, 02, and 03 to reference CALR WT sequence. Successful alignment and mismatch are shown in green and red, respectively. **D.** and **E.** Exclusion of possible off-target effects caused by CRISPR-Cas9 gene editing for *CALR* mutations. List of off-targets was provided by IDT Systems. To verify off-target sites, regions of interest were amplified by PCR and Sanger sequenced. Sequence alignment of maternal homozygous *CALR*ins5 clone (**D**) and resulting $WT_{cr}$ clones 01 and 02 for possible off-targets in *GLOD5*, *IGSF5,* and *IGSF10*. Regions of interest are highlighted in orange/grey. Sequence alignment of maternal homozygous *CALR*del52 clone (**E**) and resulting $WT_{cr}$ clones 01, 02, and 03 for possible off-targets in *CANX*, *LOC105371816, GRIN2B, RABL6,* and *BASP1P1*. Regions of interest are highlighted in orange/grey.

**Figure S2:**

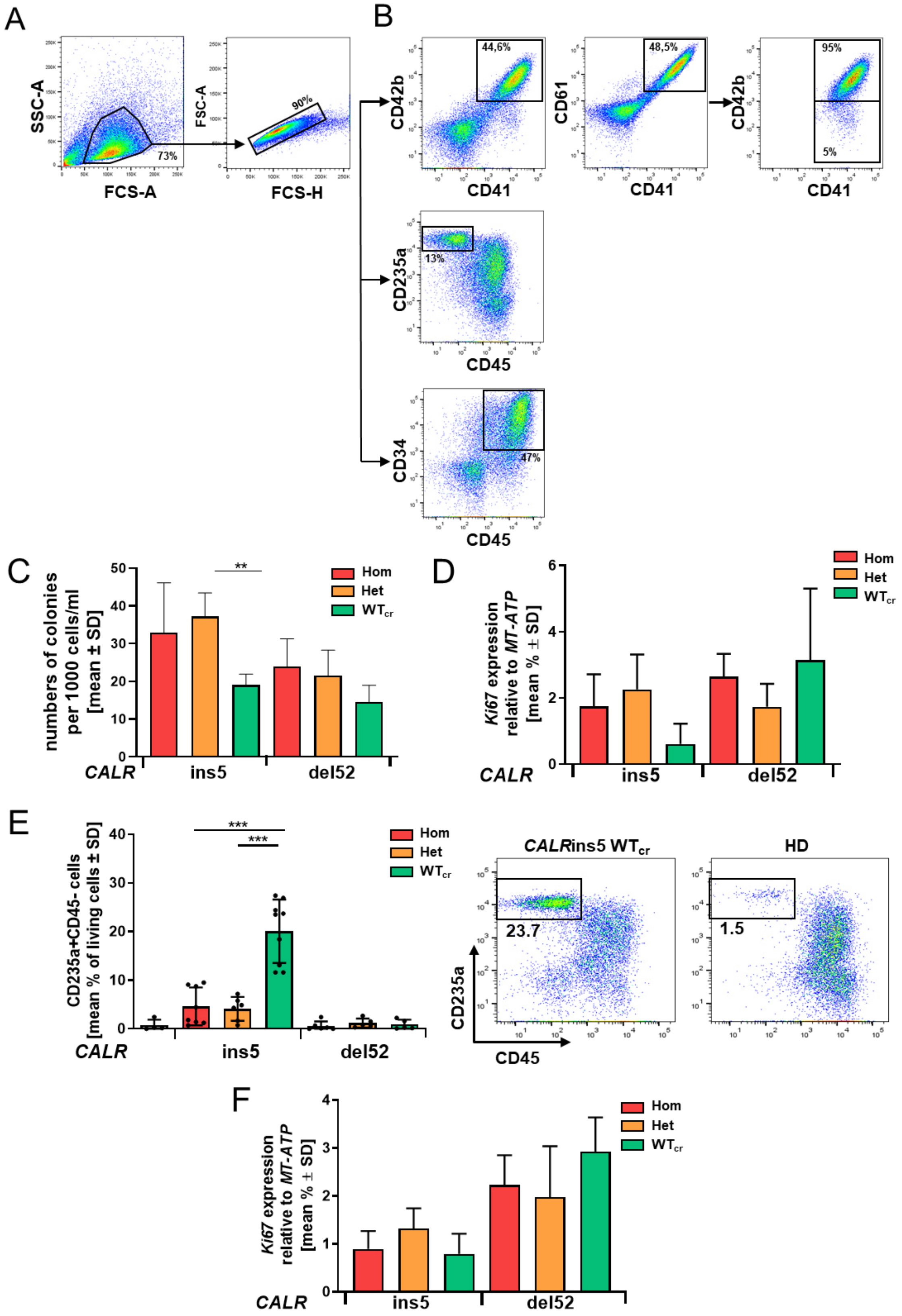
A
SSC-A
FCS-A
73%
FSC-A
FCS-H
90%
B
CD42b
CD41
44,6%
CD61
CD41
48,5%
CD42b
CD41
95%
5%
CD235a
CD45
13%
CD34
CD45
47%
C
numbers of colonies per 1000 cells/ml [mean ± SD]
**
Hom
Het
WTcr
CALR
ins5
del52
D
Ki67 expression relative to MT-ATP [mean % ± SD]
Hom
Het
WTcr
CALR
ins5
del52
E
CD235a+CD45- cells [mean % of living cells ± SD]
***
***
Hom
Het
WTcr
CALR
ins5
del52
CALRins5 WTcr
HD
23.7
1.5
CD235a
CD45
F
Ki67 expression relative to MT-ATP [mean % ± SD]
Hom
Het
WTcr
CALR
ins5
del52

**Figure S2. Impact of CALR mutations on HSCs and MKs proliferation capacity and on erythrocytic cell development**

**A.** Flow cytometry analysis gating strategy to identify myeloid cell populations in „spin-EB“ differentiation protocol. Living cells were first gated on forwards scatter (FSC-A)/side scatter (SSC-A) and further gated on single cells. **B.** Mature and immature MKs were identified by CD42b +CD41+ and CD61+CD41+, respectively. Erythroid cells were discriminated by CD235a+ CD45- expression and HSCs were determined by CD34+CD45+ expression. Numbers represent frequencies of indicated populations in percentage of living cells. **C.** Number of colonies counted from colony forming unit assay. Statistics was calculated by comparing mutated cells to corresponding $WT_{cr}$ cells. ***p*>0.01.**D.** Gene expression of the proliferative marker *Ki67* measured in iPS-derived HSC in RT-qPCR. Data are shown for three independent experiments with indicated CALR genotype. Gene expression was normalized to *MT-ATP6* expression. **E.** Gene expression of the proliferative marker *Ki67* measured in iPS-derived MKs in RT-qPCR. Data are shown for three independent experiments with indicated CALR genotype. Gene expression was normalized to *MT-ATP6* expression. **F.** Percentage of CD235a+CD45- erythroid cells determined by flow cytometry on day 14 of “spin-EB” differentiation. Number of independent experiments performed for each *CALR* genotype and HD control referred to number of data points shown. Flow cytometry plots to determine erythroid cell population (CD235a+CD45-) are exemplarily shown for *CALR*ins5 $WT_{cr}$ cells and healthy controls. Numbers represent frequencies of indicated populations in percentage of living cells. ****p*<0.001.

**Figure S3:**

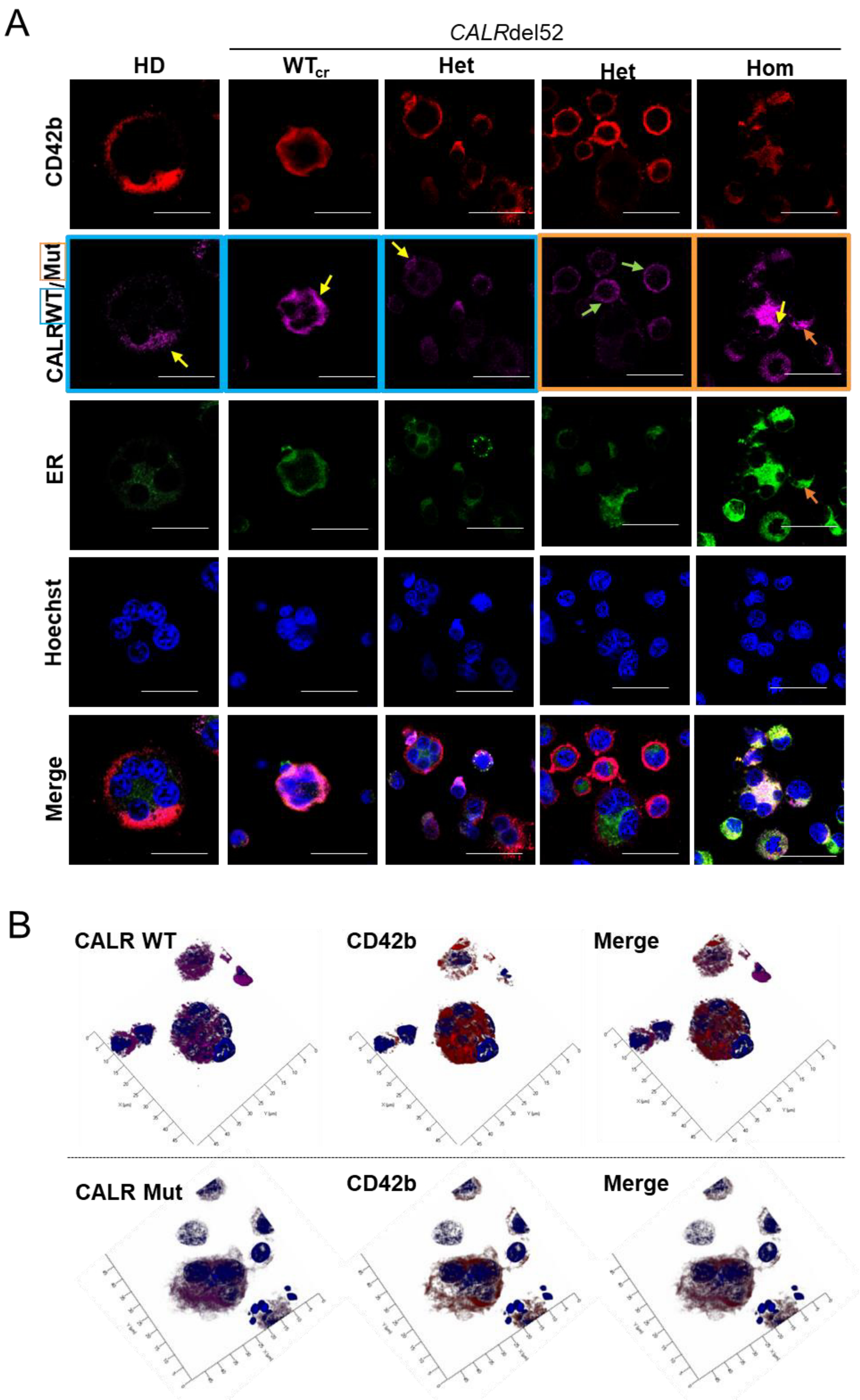
A
CALRdel52
HD
WTcr
Het
Het
Hom
CD42b
CALR WT/Mut
ER
Hoechst
Merge
B
CALR WT
CD42b
Merge
CALR Mut
CD42b
Merge

**Figure S3. Immunofluorescence staining to determine wild type and mutant CALR distribution in the cell**

**A.** iPS cell-derived MKs were fixed and stained for the endoplasmic reticulum (ER) and wild type (WT) CALR or mutated (Mut) CALR, indicated with blue or orange frames, respectively, after 14 days of differentiation for indicated iPS cell clones. To identify MKs, samples were additionally stained for CD42b. Hoechst was added for nuclear staining. Scale bars, 50 µm. Diffuse CALR distribution, clustered localization of CALR at the cell surface, and co-localization of CALR and ER are indicated with yellow, green, and orange arrows, respectively. **B.** Z-stack images of heterozygous *CALR*ins5-mutated MKs for indicated staining. Hoechst was added for nuclear staining. Scale bars, 50 µm.

**Figure S4:**

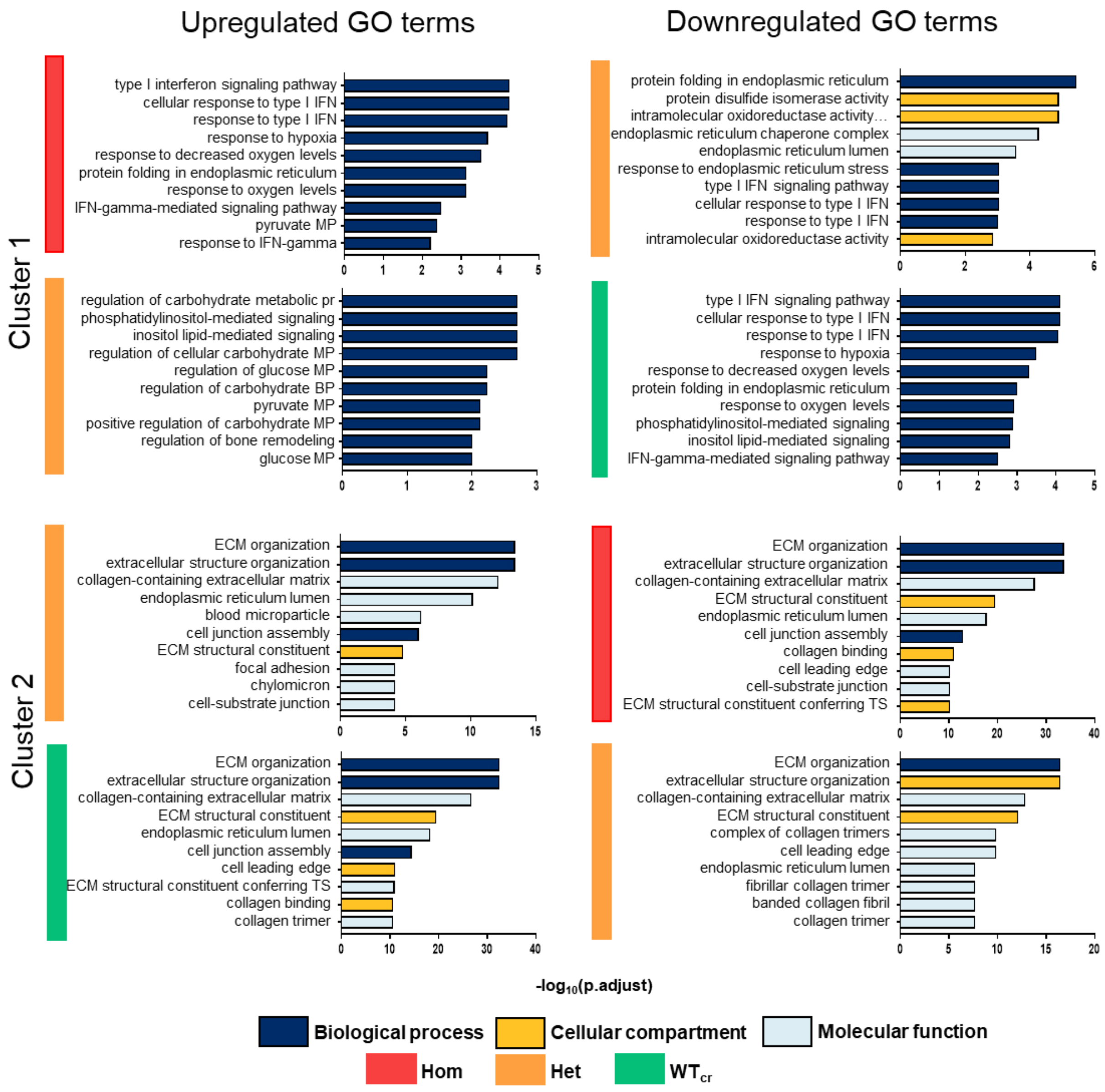

Upregulated GO terms
Downregulated GO terms
Cluster 1
Cluster 2
type I interferon signaling pathway
cellular response to type I IFN
response to type I IFN
response to hypoxia
response to decreased oxygen levels
protein folding in endoplasmic reticulum
response to oxygen levels
IFN-gamma-mediated signaling pathway
pyruvate MP
response to IFN-gamma
protein disulfide isomerase activity
intramolecular oxidoreductase activity…
endoplasmic reticulum chaperone complex
endoplasmic reticulum lumen
response to endoplasmic reticulum stress
type I IFN signaling pathway
intramolecular oxidoreductase activity
regulation of carbohydrate metabolic pr
phosphatidylinositol-mediated signaling
inositol lipid-mediated signaling
regulation of cellular carbohydrate MP
regulation of glucose MP
regulation of carbohydrate BP
positive regulation of carbohydrate MP
regulation of bone remodeling
glucose MP
ECM organization
extracellular structure organization
collagen-containing extracellular matrix
blood microparticle
cell junction assembly
ECM structural constituent
focal adhesion
chylomicron
cell-substrate junction
collagen binding
cell leading edge
ECM structural constituent conferring TS
collagen trimer
complex of collagen trimers
fibrillar collagen trimer
banded collagen fibril
-log10(p.adjust)
Biological process
Cellular compartment
Molecular function
Hom
Het
WTcr

**Figure S4. Gene ontology (GO) analysis of *CALR*ins5-mutated iPS cell derived-MKs**

Gene ontology (GO) analysis based on up- and downregulated genes of cluster-wise analysis for the heat map (Figure 6a) shown in the main text. The top 10 GO terms are shown. GO categories biological process, cellular compartment and molecular functions are depicted in dark blue, yellow, and light blue, respectively. Corresponding genotypes are shown in red, orange and green for MKs with homozygous (Hom) or heterozygous (Het) *CALR*ins5, or WTcr MKs, respectively. BP (biosynthetic process), ECM (extracellular matrix), MP (metabolic process), TS (tensile strength), IFN (interferon).

**Figure S5**

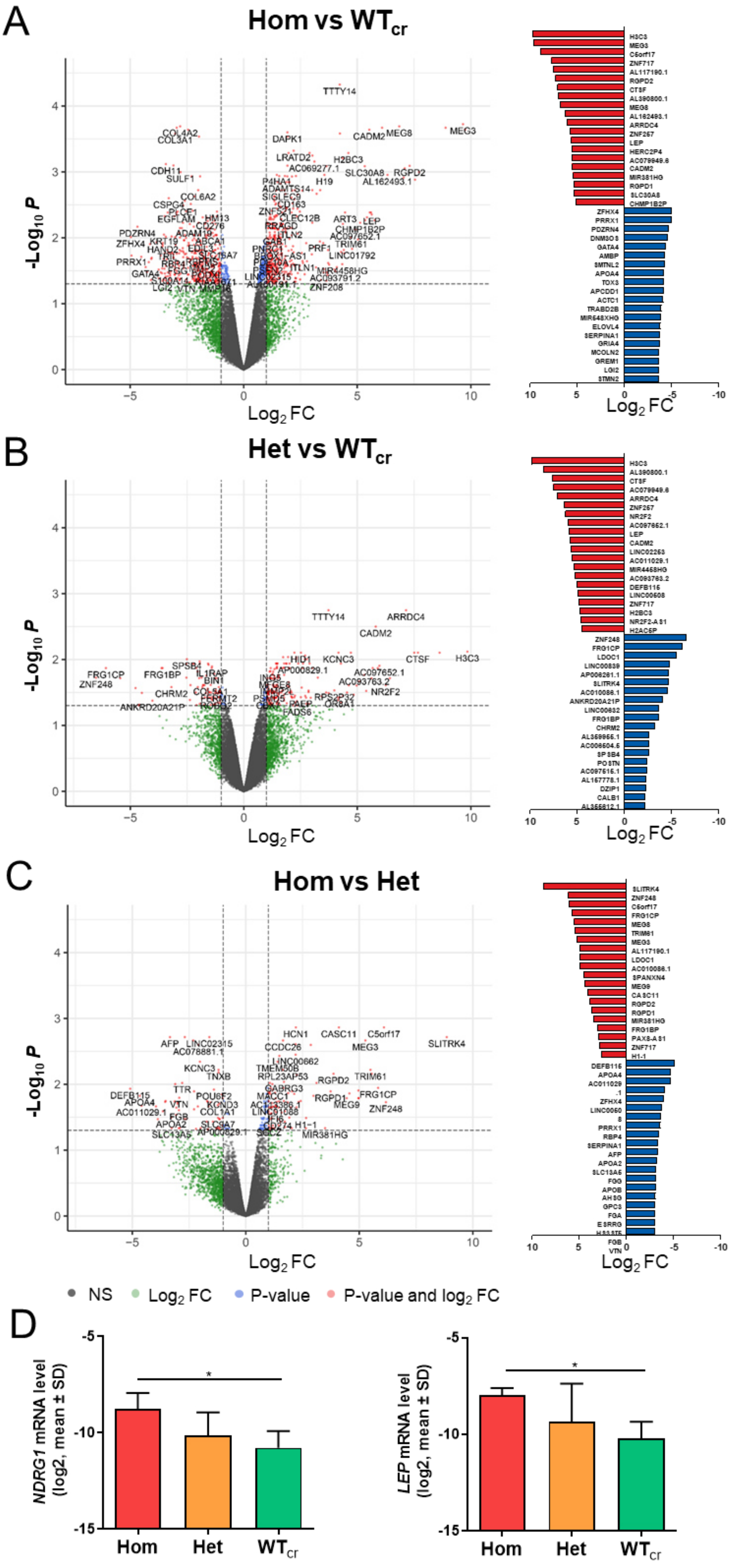
A
Hom vs WTcr
B
Het vs WTcr
C
Hom vs Het
-Log10 P
Log2 FC
NS
Log2 FC
P-value
P-value and log2 FC
D
NDRG1 mRNA level (log2, mean ± SD)
LEP mRNA level (log2, mean ± SD)
Hom
Het
WTcr
*

**Figure S5. Differentially expressed genes (DEGs) in *CALR*ins5-mutated MKs and $WT_{cr}$ MKs**

**A.**, **B.**, and **C**. Volcano plot of differentially expressed genes (DEGs) and list of the top 20 up- and downregulated DEGs among the comparisons: Hom vs $WT_{cr}$ (**A**), Het vs $WT_{cr}$ (**B**), and Hom vs Het (**C**) of CALRins5-muated MKs and control. All genes shown in the list of DEGs are significantly regulated ($p \leq 0.05$) Up-and downregulated genes are shown in red and blue, respectively. FC (fold change), Hom (homozygously *CALR*ins5-mutated), Het (heterozygously *CALR*ins5-mutated). **D.** Validation of mRNA expression of *NDRG1* and leptin (*LEP*) from RNAseq analysis by RT-qPCR isolated from iPS cell-derived MKs with *CALR*ins5 mutation or WT CALR ($WT_{cr}$) as control. RNAseq was performed using three independent experiments. RT-qPCR was performed for 2 samples also analyzed in RNAseq and 1 sample from an additional independent experiment. *$p<0.05$. Hom (homozygously *CALR*ins5-mutated), Het (heterozygously *CALR*ins5-mutated).

### SUPPLEMENTAL TABLES

#### Table S1. Generation of MPN patient-derived CALR iPS cells

| Pat. | Diagnosis | CALR mutation | Sample type | Allele burden | iPS clones screened | CALR non-mut clones | CALR Heterozygous clones | CALR homozygous clones |
|---|---|---|---|---|---|---|---|---|
| 1 | pET-MF | ins5 | PB | 82% | 101 | 0 | 27 | 74 |
| 2 | PMF | del52 | PB | 51% | 64 | 0 | 55 | 9 |
| 3 | PMF | del31 | PB | 37% | 69 | 5 | 64 | 0 |

**Table S 1.** iPS cells were generated by reprogramming peripheral blood-derived mononuclear cells from three patients carrying *CALR* del52, ins5, or del31 mutation. Individual colonies were picked and screened for *CALR* genotypes by PCR. Abbreviations: CALR, calreticulin; iPSC, induced pluripotent stem cell; non-mut, non-mutant; PMF, primary myelofibrosis; pET-MF, secondary post essential thrombocythemia myelofibrosis. PB, peripheral blood.

#### Table S2. CRISPR guide RNAs (gRNAs) to repair homozygous CALR mutations with CRISPR/Cas9

| Mutation | gRNA |
|---|---|
| *CALR*ins5 | 5'-AGGCAGAGGACAATTGTCGG -3' |
| *CALR*del52 | 5'-ACGAGGAGCAGAGGACAAGG-3' |

#### Table S3. Donor template to repair homozygous CALR mutations with CRISPR/Cas9

| Mutation | Donor Template |
|---|---|
| *CALR*ins5 | 5'-GAGGCTTAAGGAGGAGGAAGAAGACAAGAAACGCAAAGAGGAGGAGGAGGCAGAGGACAAGGAGGATGATGAGGACAAAGATGAGGATGAGGAGGATGAGGAGGACAAGGAGGAAGATGA -3' |
| *CALR*del52 | 5'-GTGCTCTGCCTGCAGGCAGCAGAGAAACAAATGAAGGACAAACAGGACGAGGAGCAGAGGCTTAAGGAGGAGGAAGAAGACAAGAAACGCAAAGAGGAGGAGGAGGCAGAGGACAAGGAGGATGATGAGGACAAAGATGAGGATGAGGAGGATGAGGAGGACAAGGAGGAAG -3' |

### Table S4. NGS Data of patient-derived CALR iPS cells and Healthy donor

| | clone | HD | | *CALR* ins5 (K385fs) | | | | | | *CALR* del52 (L367fs) | | | | | | *CALR* del31 (E372fs) | | | |
|---|---|---|---|---|---|---|---|---|---|---|---|---|---|---|---|---|---|---|---|
| | | HD 4 | HD 6 | WT16 | WT18 | Het04 | Het10 | Hom13 | Hom49 | WT27 | WT30* | WT44 | Het01 | Hom34 | Hom41 | non-mut41 | non-mut48 | Het06 | Het24 |
| Mutation | *CALR* ins5 | | | | | 43 | 41 | 85 | 93 | | | | | | | | | | |
| | *CALR* del52 | | | | | | | | | | | | 60 | 97 | 97 | | | | |
| | *CALR* del31 | | | | | | | | | | | | | | | | | 54 | 55 |
| Polymorphism | JAK2 R1063H | 51 | | | | | | | | 46 | 48 | 48 | 52 | 47 | 48 | | | | |
| | JAK2 S518F | | | | | | 5 | | | | | | | | | | | | |
| | TET2 L1721W | 50 | | | | | | | | 50 | 51 | 51 | 49 | 49 | 46 | 50 | 47 | 48 | 50 |
| | TP53 P72R | 59 | | 100 | 99 | 100 | 100 | 100 | 99 | 63 | 51 | 55 | 55 | 63 | 46 | 100 | 99 | 100 | 99 |
| | ASXL1 L815P | | | 100 | 100 | 100 | 100 | 100 | 100 | | 100 | 100 | | | | | 100 | | 100 |
| | ETNK1 Q284R | | | 46 | 47 | 50 | 53 | 51 | 48 | | | | | | | | | | |
| | KIT M541L | | | 43 | 48 | 47 | 41 | 51 | 53 | | | | | | | | | | |
| | U2AF1 5'UTR | | | | | 55 | 56 | 52 | | | | | | | | | 53 | | 53 |
| | CBL M63I | | | | | | 6 | | | | | | | | | | | | |
| | MPL L512P | | | | | | 6 | | | | | | | | | | | | |
| | SETBP1 V231L | | | | | | | | | | 48 | | | | | | | | |
| | SETBP1 V1101L | | | | | | | | | | 50 | | | | | | | | |
| | SH2B3 W262R | | | | | | | | | | 100 | | | | | | | | |
| | BARD1 C557S | | | | | | | | | | | | | | | 52 | 50 | 52 | 49 |

**Table S4.** Next generation sequencing (NGS) of patient-derived iPS cell clones and healthy donors iPS cell clones. All identified clinically relevant mutations and polymorphisms are listed. Percentages represent allele frequency (the fraction of mutant reads per total reads) of indicated mutations or polymorphisms.

**Table S5: List of antibodies used for flow cytometry**

| **Antibody** | **Company** |
|---|---|
| APC mouse anti-human CD34 | BD, USA |
| FITC mouse anti-human CD43 | BD, USA |
| PE mouse anti-human CD31 | BD, USA |
| FITC anti-human CD61 clone VI-PL2 | BioLegend, USA |
| Anti-human CD117 (ckit)-PE Cy7 clone 104D2 | Thermo Fisher Scientific, USA |
| Anti-human CD235a-PE clone HIR2 | Thermo Fisher Scientific, USA |
| CD45-APC-Vio770 human | Miltenyi Biotec, Germany |
| CD66b-PE human | Miltenyi Biotec, Germany |
| APC anti-human CD42b | BioLegend, USA |
| PE/Cyanina7 anti-human CD41 | BioLegend, USA |
| CD45-APC human | Beckman Coulter. USA |
| CD15-PE Cy5 human | Beckman Coulter, USA |
| MPO-FITC human | Beckman Coulter, USA |

**Table S6: List of primary and secondary antibodies used for Western blot and Immuofluorescence**

| **Antibody** | **Company** |
|---|---|
| Monoclonal anti-mutated Calreticulin Rabbit | Dianova, Germany |
| Monoclonal anti Calreticulin (D3E6) XP Rabbit #12238 | Cell Signaling, USA |
| Monoclonal GAPDH (0411): sc-32233 Mouse | Santa Cruz Biotechnology, USA |
| CD42b Polyclonal Antibody | Thermo Fisher Scientific, USA |
| Calreticulin Antibody (1G6A7) | Novus Biologicals, USA |
| Goat anti-rabbit IgG (H+L) Alexa Fluor 555 | Thermo Fisher Scientific, USA |
| Goat anti-mouse IgG (H+L) Alexa Fluor 647 | Thermo Fisher Scientific, USA |
| Goat anti-rabbit IgG (H+L) FITC | Thermo Fisher Scientific, USA |
| Goat anti-mouse IgM (H+L) Alexa Fluor 594 | Thermo Fisher Scientific, USA |
| Goat anti-mouse Immunoglobulins/HRP, polyclonal | Dako, USA |
| Goat anti-rabbit Immunoglobulins/HRP, polyclonal | Dako, USA |

**Table S7: List of primers used for qPCR and CALR genotyping PCR**

| Target name | | Sequence | Reference |
|---|---|---|---|
| **qPCR** | | | |
| *CALR*del52 | Non-.mut allele FRW | CAGGACGAGGAGCAGAGACT | |
| | Mutant allele FRW | ACAGGACGAGGAGCAGAGAAC | |
| | Common Rev | GCCTCTCTACAGCTCGTCCTTG | |
| *CALR*del31 | Non-.mut allele FRW | CAAGTCTGGCACCATCTTTG | |
| | Mutant allele FRW | TCCTCTTTGCGTTTCTTGTC | |
| | Common Rev | ATCCTCCTTGTCCTCTGTTC | |
| *CALR*ins5 | common FRW | CAAGTCTGGCACCATCTTTG | |
| | Non-mut allele Rev | TGTCCTCATCATCCTCCTTG | |
| | mutant Rev | TGTCCTCATCATCCTCCGAC | |
| *ETS1* | FRW | TCGATCTCAAGCCGACTCTC | |
| | REV | CATTCACAGCCCACATCACC | |
| *FLI1* | FRW | GTGCTGTTGTCACACCTCAG | |
| | REV | TACTGATCGTTTGTGCCCCT | |
| *GAPDH* | FRW | GTTGAGGTCAATGAAGGGGTC | |
| | REV | GACCTCAACTACATGGTGAGTTGC | |
| *GATA1* | FRW | GGGATCACACTGAGCTTGC | [1] |
| | REV | ACCCCTGATTCTGGTGTGG | |
| *GFI1B* | FRW | AGTTCTGCGGCAAGCGTTTCCA | |
| | REV | TTTCCGCACACCTGGCACTTGT | |
| *Ki67* | FRW | CAGACCCATTTACTTGTGTTGGA | |
| | REV | ACGCCTGGTTACTATCAAAAGG | |
| *LEP* | FRW | CGGTAAGGAGAGTATGCGGG | |
| | REV | ACCAGAAAGAGTGGAGCCT | |
| *LOX1* | FRW | GCATACAGGGCAGATGTCAGA | |
| | REV | TTGGCATCAAGCAGGTCATAG | |
| *MPL* | FRW | CTGCCACTTCAAGTCACGAA | |
| | REV | CTGCCACTCCAATTCCAGAT | |
| *MPO* | FRW | CCGGGATGGTGATCGGTTTT | [2] |
| | REV | CAGATGATCCGGGGCAATGA | |
| *MT-ATP6* | FRW | CGTACGCCTAACCGCTAACA | |
| | REV | AGGCGACAGCGATTTCTAGG | |
| *NRDG1* | FRW | GACAAAGGCCAAAAGGTCAACA | |
| | REV | CCATTTCATTGGGAGGGTGGT | |
| *NFE2* | FRW | CTGTGACTCCACCACAGGTTT | |
| | REV | TGAGCAGGGGCAGTAAGTTG | |
| *RUNX1* | FRW | CCGAGAACCTCGAAGACATC | |
| | REV | GTCTGACCCTCATGGCTGT | |
| *vWF* | FRW | CAACACCTGCATTTGCCGAA | |
| | REV | TGACCTGTGACAAGGCACTC | |
| **Genotyping PCR** | | | |
| *CALR*del52 r | FRW | ACAACTTCCTCATCACCAACG | |
| *CALR*del31 | REV | GGCCTCAGTCCAGCCCTG | |
| **Gentoyping allele specific PCR** | | | |

| | | |
|---|---|---|
| *CALR*ins5 | common FRW | TAACTGCAGTGTCAGCGGTG |
| | Non-mut allele Rev | TGTCCTCATCATCCTCCTTG |
| | mutant Rev | TGTCCTCATCATCCTCCGAC |
| **CRISPR-off target products PCR** | | |
| *CANX* | FRW | ACACGTCTTCAGGGTAGGA |
| | REV | CAACATCGTAGGGTCTTGGCT |
| *RABL6* | FRW | GCAAAGAGGTACTGGCTACTCC |
| | REV | CGGCCTAGAGCTCCTCGTA |
| *BASP1P1* | FRW | GGCGGAGCTAGCACTACAAC |
| | REV | GGTGACTTCGGCAGCTTTGG |
| *IGSF10* | FRW | AAGTGAGTGAACCCAGGCAC |
| | REV | AGCTTTGGGGAAGGTGATGG |
| *IGSF5* | FRW | TAGAGATTCCTGGTTCCTGGG |
| | REV | TCCACCGCCACGTCCTAGATT |
| *GRIN2B* | FRW | TCACCACACACGCTACTTCCAC |
| | REV | TTTACAGAGAAGGCTGGCCG |
| *GLOD5* | FRW | CCTTCCATTTGCACTACCTACCT |
| | REV | CCCTGGCTAAACTGGGGGAG |
| *LOC105371816* | FRW | TCTTGGAGACAGACTGCTGG |
| | REV | GTGTGGGGTCCTGATGCTTTA |

**Table S8: Medium composition**

| **SFM Medium for “spin-EB” differentiation** | | |
|---|---|---|
| 50 % | IMDM | Thermo Fisher Scientific, USA |
| 50 % | Ham’s F-12 Nutrient Mixture | Thermo Fisher Scientific, USA |
| 0.5 % | Human Plasbumin 20 | Grifols, Germany |
| 2 mM | GlutaMAX | Thermo Fisher Scientific, USA |
| 2 mM | Chemically Defined Lipid Concentrate | Thermo Fisher Scientific, USA |
| 50 µg/ml | L-ascorbic acid | Stemcell Technologies, Canada |
| 6 µg/ml | Transferrin | Merck, Germany |
| 400 µM | 1-Thioglycerol | Merck, Germany |

| **EB medium for EB-based differentiation** | | |
|---|---|---|
| | IMDM | Thermo Fisher Scientific, USA |
| 15 % | FBS | Pan Biotech, Germany |
| 1U/ml | Penicillin | Thermo Fisher Scientific, USA |
| 100 mg/ml | Streptomycin | Thermo Fisher Scientific, USA |
| 2 mM | L-Glutamine | Thermo Fisher Scientific, USA |
| 5 % | Protein-Free Hybridoma Medium-II | Thermo Fisher Scientific, USA |
| 0.1 mM | β-mercaptoethanol | Thermo Fisher Scientific, USA |
| 50 µg/ml | L-Ascorbic acid | Sigma Aldrich, Germany |
| 175 µg/ml | hTransferrin | Sigma Aldrich, Germany |

| Progenitor medium for EB-based differentiation | | |
|---|---|---|
| | StemPro 34 SFM | Thermo Fisher Scientific, USA |
| 2 mM | L-Glutamine | Thermo Fisher Scientific, USA |
| 1U/ml | Penicillin | Thermo Fisher Scientific, USA |
| 100 mg/ml | Streptomycin | Thermo Fisher Scientific, USA |
| 1X | MEM Non-Essential Amino Acids | Thermo Fisher Scientific, USA |
